# Warming and biotic interactions shape selection on plant traits and their plasticity

**DOI:** 10.64898/2026.09.28.754985

**Authors:** Juan Traine, Florian P. Schiestl, Quint Rusman

## Abstract

- Global warming will impact natural selection on plant traits and their plasticity by influencing phenotypes, biotic interactions, and fitness. Because warming affects biotic interactions, community composition may influence how warming affects selection on plants, but this is little studied.
- We grew full-sibling pairs of *Brassica rapa* plants in an ambient and hot temperature environment and exposed them to visitation by a pollinator (*Bombus terrestris*), a pollinating herbivore (*Pieris rapae*), or their combination. We recorded plant phenotypes, insect visitation, herbivory, and plant fitness. We used these to calculated phenotypic selection on plant traits and their plasticity, including additive and nonadditive effects.
- Elevated temperature changed selection on traits related to pollinator attraction, such as flower number, nectar production, and floral scent emission. We found both positive and negative selection on temperature-mediated trait plasticity depending on the biotic treatment. Elevated temperature changed patterns of additive and non-additive selection on a variety of traits including flower morphology and scent.
- This study shows that elevated temperatures can reshape patterns of phenotypic selection, including nonadditive effects, and plastic trait expression. Effects of elevated temperature on phenotypic selection depended on the biotic interactors, highlighting the importance of community context for the effects of warming on natural selection.

## Introduction

Global warming is driving an increase in average temperatures and extreme temperature events, and such rapid environmental shifts may impact the survival and reproduction of organisms. Thereby, global warming potentially influences patterns of phenotypic selection (Jump *et al*., 2006; Campbell & Powers, 2015). Phenotypic selection occurs when individuals with certain phenotypes differ in survival or reproduction compared to individuals with other phenotypes, and can be measured as trait-fitness relationships (Lande & Arnold, 1983; Kingsolver & Pfennig, 2007). For plants, warming can impact selection directly by affecting plant physiology and fertility (Zhang *et al*., 2017; Borghi *et al*., 2019; Traine *et al*., 2024). Warming can also modify the performance and behaviour of biotic selection agents such as herbivores and pollinators (Scaven & Rafferty, 2013; Hamann *et al*., 2021; Rusman *et al*., 2025b). For example, under experimental warming, solitary bees spent less time on flowers and were less specialized in their visitation, with negative consequences for plant fitness (de Manincor *et al*., 2023). Thus, by directly affecting both plants and their biotic interactors, warming may lead to changes in phenotypic selection with potential consequences for plant evolutionary trajectories (Becklin *et al*., 2016; Anderson & Song, 2020; Rusman *et al*., 2025b).

Temperature can further impact selection on plant traits through temperature-mediated phenotypic plasticity (Dorey & Schiestl, 2022; Gómez *et al*., 2024; Traine *et al*., 2024). Plants display various plastic responses which help them deal with increasing temperatures. For example, reductions in flower size and lighter colours can reduce the amount of radiant energy absorbed by flowers and thus help regulate the temperature under hotter conditions (van der Kooi *et al*., 2019). Such changes in trait expression with potential consequences for plant fitness may change trait-fitness relationships and thereby alter selection (Hunter *et al*., 2018). Moreover, temperature-induced plant responses can impact how plants interact with their biotic environment, with potential consequences for biotic-mediated selection (Gérard *et al*., 2020; Hamann *et al*., 2021). For example, temperature-mediated floral plasticity altered pollinator visitation, plant fitness, and pollinator-mediated selection (Gómez *et al*., 2020; Descamps *et al*., 2021; Traine *et al*., 2024; Gómez *et al*., 2025). When temperature-induced plant responses provide a fitness benefit, such plasticity can be under selection to be maintained or increased (Napier *et al*., 2023; Gómez *et al*., 2024). However, plasticity can be under negative selection when it has costs, such as physiological costs to maintain plastic responses or ecological costs such as reduced pollinator attraction (Van Kleunen & Fischer, 2005; Gómez *et al*., 2024). Understanding how temperature and biotic agents interact to shape selection on temperature-induced plasticity will help understand if such plant responses can help plants adapt to global warming.

Changes in biotic communities under global warming will alter patterns of phenotypic selection in more complex ways. Selection in multispecies communities is diffuse and depends on community context (Janzen, 1980; Strauss & Irwin, 2004). In other words, the strength of selection on traits changes depending on which selection agents are present (Strauss *et al*., 2005; Sahli & Conner, 2011). Diffuse selection by multiple species can be additive, the sum of individual pairwise selection gradients of each species, or nonadditive, where combined selection is not equal to the sum of individual pairwise selection gradients (Strauss *et al*., 2005; Haloin & Strauss, 2008; terHorst *et al*., 2018). For example, selection by butterflies and bumblebees on floral scent of *Brassica rapa* was higher or lower for scent compounds than expected from selection by butterflies or bumblebees alone (Knauer & Schiestl, 2017). Nonadditive selection is caused by genetic correlations between traits or because the total strength of selection on a trait is altered by another interacting species (Strauss *et al*., 2005; terHorst *et al*., 2018). Because global warming can alter the identity of selection agents (Zhu *et al*., 2015; Toftegaard *et al*., 2019), the sign and strength of selection of each individual selection agent (Traine *et al*., 2024; Rusman *et al*., 2025b), and interactive effects between selection agents, it is highly likely that global warming will alter patterns of nonadditive selection. Investigating the effects of elevated temperatures on diffuse selection will improve our understanding of how environmental change will alter selective landscapes.

Our previous work investigated how warming, when solely affecting plants, could impact plant-insect interactions, plant fitness and selection through temperature-mediated plant plasticity (Traine *et al*., 2024). In this study, both plants and insects were exposed to warming. The current study was part of a larger experiment to investigate how warming influenced plant evolution in a community context (Rusman *et al*., 2025a; Traine *et al*., 2026a; Traine *et al*., 2026b). Therefore, the current study overlaps experimentally with a study on the effects of warming on reciprocal selection between plants and butterflies (Rusman *et al*., 2025b). Here, we aimed to determine whether warming, through direct and indirect effects on both plants and insects, could impact 1) fitness and traits under selection of *B. rapa* plants, 2) selection on temperature-mediated trait plasticity, and 3) complex patterns of diffuse selection by a pollinator, *Bombus terrestris*, and a pollinating herbivore, *Pieris rapae*. We expected warming to negatively impact plant fitness and change natural selection on plant traits due to plastic reductions in most floral traits and subsequent negative impacts on pollinator visitation (Traine *et al*., 2024; Rusman *et al*., 2025b). We expected temperature-mediated phenotypic plasticity of floral advertisement and reward traits to be under negative selection to reduce ecological costs, *e.g.* reduced pollinator visitation (Descamps *et al*., 2021; Gómez *et al*., 2024; Traine *et al*., 2024).

## Methods

### Plant and insects

We used fast-cycling *Brassica rapa* plants (Wisconsin Fast Plant, Carolina Biological Supply, Burlington). *Brassica rapa* is an annual/biannual, mostly self-incompatible plant with a diverse pollinator community including cabbage white butterflies (*Pieris rapae*) and buff-tailed bumblebees (*Bombus cf. terrestris*) (Atmowidi *et al*., 2007; Rader *et al*., 2009), and a diverse herbivore community that includes *P. rapae* caterpillars (Agren & Schemske, 1993). We grew 400 plants, paired them randomly, and cross-pollinated them by hand. The seeds from both parents within each pair were collected and became full-sib families (groups of seeds that share both parents) (Traine *et al*., 2024; Rusman *et al*., 2025b). We used 98 full-sib families in this study.

Of each full-sib family, 12-16 seeds were sown in four cohorts which one week between each cohort of 3-4 seeds. Seeds were sown in multi-pot trays filled with a sowing mix consisting of equal parts sterilized sieved topsoil (Zurich Botanical Garden, Switzerland), compost (3 years-old, Zurich Botanical Garden, Switzerland), pumice (1-5 mm, Ökohum GmbH, Switzerland), and peat (medium structure, Zelta Zeme, Latvia). We germinated the seeds for 8 days in a phytotron (24hs light, 22±1°C, 60%RH). On day 8 we re-potted seedlings individually (588 in total, 6 per full-sib family spread over 4 cohorts), in pots filled with standardized soil (Einheitserde, Patzer-Erde GmbH, Sinntal, Germany). After re-potting, we assigned half the plants to a greenhouse cabin with an ambient temperature treatment (23 C) and the other half to a greenhouse cabin with a hot temperature treatment (27 C, plus six weekly 30°C days). These temperature parameters approximate the current and predicted mid-century mean early summer temperatures during the day in northern Switzerland (Fischer *et al*., 2022). Even though we used one greenhouse cabin per temperature treatment, we show with an experiment that the greenhouse cabins and temperature treatment were independent, and the effects of our temperature treatment were caused by differences in temperature rather than differences between the greenhouse cabins (see supplementary information “*Experiment 2”*, Table S1,S2). Plants were watered twice a day, morning and afternoon, and supplementally watered during the day if too dry.

We used *Bombus terrestris* bumblebees as pollinators, and *Pieris rapae* butterflies as pollinating herbivores (mutualist/antagonist), since they feed on nectar and pollinate *B. rapa*, but also oviposit on plants and caterpillars develop successfully on fast-cycling *B. rapa* (Traine *et al*., 2024; Rusman *et al*., 2025b). See supplementary methods for rearing conditions and origins.

### Flowering plant phenotyping

To investigate the effects of warming on the flowering phenotype of *B. rapa* plants, we monitored plant flowering phenology, and measured morphology, flower volatile emission, and nectar production 21-25 days after sowing. Detailed descriptions of trait measurements can be found in the supplementary methods.

### Bioassays

To study the effects of warming on biotic interactions, we performed pollination and herbivory bioassays at ambient and hot conditions. At both temperatures, we exposed plants to pollination by bumblebees (*Bombus-*treated plants), pollination and herbivory by butterflies/caterpillars (*Pieris-*treated plants) or pollination by bumblebees and pollination/herbivory by butterflies/caterpillars (generalized-treated plants), resulting in six treatments. Each treatment had 98 plants divided into two replicates (A and B) containing 49 plants. Each plant came from a different full-sib family, and the same families were used across treatments. Due to timing issues with butterfly availability, *Bombus*-treated plants comprised seed cohort 1 (rep. A) and 2 (rep. B), while *Pieris-* and generalized-treated plants comprised seed cohort 3 (rep. A) and 4 (rep. B). Bioassays were conducted 24-27 days after plant sowing, and always after trait measurements had been done. Each replicate of 49 plants was arranged in a 7×7 matrix inside a flight cage (2.5×1.8×1.2 m).

For *Bombus*-treated plants, we released 12 individual worker bumblebees sequentially. Each bee could visit 3 plants before re-capture. We recorded which plants were visited, the number of flower visits per plant, and the time (in seconds) spent per plant. We divided the time spent per plant by the number of flower visits to calculate the time spent per flower (in seconds).

For *Pieris*-treated plants, we released a total of 25 mated female butterflies for 4 days to visit flowers and oviposit on plants (Rusman *et al*., 2025b). Butterflies were mated 2 days before the assays, starved for 24 hs before, and had no contact with any plants before the assays. On the first two days, 13 butterflies were released, and on the last two days, 12 butterflies. Release happened at 9:00am and recapture at 17:00pm, during which we recorded each flower visit and the number of eggs laid per plant. We used a voice recorder (VoiceTracer DVT4110, Philips, Amsterdam, The Netherlands) to record our observations, which helped us keep track of the butterflies. Plants were ranked for their egg load (rank 1 having the most eggs and rank 49 having the least eggs) and assigned to five herbivory treatments based on this rank. Plants ranked 1-9 kept four caterpillars, 10-19 three caterpillars, 20-29 two caterpillars, 30-39 one caterpillar, and 40-49 zero caterpillars. We removed all excess larvae 7-9 days after oviposition. The ranking system ensured herbivory proportional to oviposition while avoiding caterpillar amounts >4 which would kill the plant (Traine *et al*., 2024; Rusman *et al*., 2025b). A few days later, we rechecked the caterpillar numbers per plant and fitted plastic reverse cones made from 3 transparent PVC A4-sized pages around each plant. These cones had an open top, with its border enveloped in cotton wool, preventing caterpillars from switching plants without hindering plant growth. When all caterpillars from each plant had pupated, we removed the cones.

For generalized-treated plants, we released butterflies and bumblebees following similar protocols as described above, but we reduced the number of insects released for each species so that total pollination would be comparable to that of the other treatments. We released 6 bumblebees and 20 butterflies simultaneously; bumblebee bioassays were performed while butterflies were inside the same flight cage. All butterflies were released simultaneously for 2 days. Observations were performed for 2 h per day, for 4hs total, and always after bumblebee bioassays. We managed butterfly oviposition and herbivory in the same way as described above.

### Plant reproductive output

One month after the bioassays, we moved the plants to a different greenhouse for drying and fruit collection. Once dry, we counted the number of undamaged fruits and fruits showing evidence of caterpillar damage. We then opened every fruit per plant, emptied the seeds on a white plate, and photographed it. Using these photographs, we counted the number of seeds per plant using the software ImageJ. We divided the number of damaged fruits by the total number of fruits to calculate the proportion of damaged fruits, the number of fruits by the number of open flowers at the time of pollination to calculate the fruit set (capped at 1), and the number of seeds by the number of fruits to calculate the number of seeds per fruit.

### Statistical methods

We analysed differences in traits of plants grown in the two temperature environments using (generalized) linear mixed models ((G)LMMs). Each trait was tested as a dependent variable against temperature (ambient and hot) as a fixed factor. Planting week (cohort), replicate and plant family were included as random factors. For floral scent compounds, the sampling date was also added as a random factor and data was log-transformed.

The dataset to analyse differences in traits of plants grown in the two temperature environments, (and used for all selection analyses), comprised all plants, including *Pieris*-treated plants. However, data from *Pieris-*treated plants used here (phenotypes, plant-butterfly interactions, reproduction) is already described in Rusman *et al*. (2025b). Still, our dataset comprised all plants including data from *Pieris-*treated plants, because we were interested in the overall effects of temperature on plant trait expression and selection patterns, and differences in the effects of temperature on selection depending on biotic treatment.

We analysed differences in insect visitation and oviposition between temperatures using (G)LMMs. We tested the effect of temperature on the number of plant and flower visits (bumblebees and butterflies, see also Rusman *et al*. (2025b)), time spent per plant and per flower (bumblebees). The number of flower visits from the generalized treatment was calculated by adding the number of visits by butterflies and bumblebees per plant. Replicate and plant family were included as random factors. To determine whether insects were attracted by different plant traits between temperature treatments, we performed generalized linear mixed models, which are described in the supplementary information.

We analysed the differences of plant reproductive output between temperature environments using GLMMs. For each biotic treatment separately, we tested the effect of temperature on fruit set, number of fruits, seeds, and seeds per fruit, and the proportion of damaged fruits (treatments including butterflies, see also Rusman *et al*. (2025b). Replicate and plant family were included as random factors.

For all traits and variables, if normality assumptions were met, we modelled continuous data using a Gaussian distribution. Non-normal continuous data was modelled using a Gamma distribution with a log link function or a Tweedie distribution if we detected zero-inflation. We used a beta (data without 0 and 1s) or ordered beta (data with 0 and 1s) distribution for proportional data. We modelled count data using a Poisson distribution, a negative binomial distribution with a log link function if data was overdispersed and/or zero-inflated. Time until flowering was analysed using a Cox proportional hazards distribution. All packages used are described in the supplementary information.

#### Effects of temperature on linear and non-linear selection

We tested if selection on plant traits differed between temperature and biotic treatments. Linear and nonlinear selection gradients were estimated using multiple regression (MR) analyses, with relative fitness as dependent variable and standardized traits as covariates (Lande & Arnold, 1983). Selection analyses on phenological/visual traits and floral scent compounds were performed separately, because visual and scent traits were generally not correlated (Dorey & Schiestl, 2022; Traine *et al*., 2024; Rusman *et al*., 2025b). For each trait group, we included all phenological/visual traits and floral scent compounds, except petal length and width (composites of petal area), and total volatile emission (sum of individual compounds). Collinearity was not a problem with most traits having a VIF of <10 (Morrissey & Ruxton, 2018). We did not regularize our MRs because classic MR models perform equally well as regularized MR models when VIF<20 while reporting unbiased selection estimates (Sztepanacz & Houle, 2024). We analysed selection within each pollinator treatment separately.

To ascertain the interactive effects of temperature on selection, we first estimated linear and nonlinear selection from MR-models including “temperature environment” as a fixed factor and its interactions with each trait. After, linear and nonlinear selection were estimated for each separate temperature environment. We obtained linear selection gradients from models including only linear terms. More details for nonlinear selection analyses are provided in the supplementary information. The MR-models for *Bombus-*treated plants did not meet normality criteria due to the many zeros because about half the plants did not get visits. Thus, estimates, standard errors, and 95% bias-corrected confidence intervals were calculated through an ordinary nonparametric bootstrap with 10 000 iterations.

We did not correct our p-values from MR-models for multiple testing. Multiple testing corrections seek to maintain the type I error for the omnibus null and not reduce the type I error for the surrogate nulls (Perneger, 1998; Frane, 2021; Rubin, 2021; García-Pérez, 2023). In our case, the omnibus null would be that the specific suite of traits included all influence fitness, while the surrogate nulls (or individual tests) test if each individual traits influence fitness. We are interested in and test the surrogate nulls, rather than the omnibus null. Since corrections for multiple testing will not reduce the type I error rate of the individual tests but rather increase the chance of type II errors (Perneger, 1998; Frane, 2021; Rubin, 2021; García-Pérez, 2023), we preferred to avoid them.

We calculated selection differentials for each trait using individual linear regressions between relative fitness (plant seed number divided by the average replicate seed number) and each standardized character (Lande & Arnold, 1983) of plants from each temperature and biotic treatment separately. Functions and packages are described in the supplementary information.

#### Selection on temperature-mediated plasticity

We tested if selection on temperature-mediated plasticity of each trait differed between biotic treatments by 1) estimating selection on trait plasticity and 2) inferring the adaptive value of plasticity (Gómez *et al*., 2024). We estimated selection on temperature-mediated plasticity by correlating trait plasticity (measured as the slope of the reaction norm between traits expressed at different temperatures) of a full-sib family with the overall fitness of the family across temperatures (Van Kleunen & Fischer, 2005). We estimated selection using MR analyses where total relative fitness per family was the dependent variable and the standardized slopes of the temperature reaction norms of each trait were the covariates (Gómez *et al*., 2024). Total relative fitness of each plant family was calculated as the sum of seeds per family pair divided by the average family sum of seeds per replicate, and for each pollination treatment separately. To obtain temperature reaction norms for each trait we calculated the difference between the trait values of sibling plants grown in the hot and ambient environment (Ramos & Schiestl, 2020; Wang *et al*., 2022). For all analyses, selection gradients, standard errors, p-values, and confidence intervals were obtained similar as described above.

We inferred the adaptive value of temperature-mediated plasticity by comparing the direction of (significant) selection in the ambient and hot environment, and the direction of plasticity (Gómez *et al*., 2024). We considered temperature-mediated plasticity adaptive when selection in the hot environment was significant and in the same direction as plasticity, while selection in the ambient environment was either non-significant or significant and in the opposite direction as plasticity. We considered temperature-mediated plasticity maladaptive when selection in the hot environment was significant and in the opposite direction of plasticity, while selection in the ambient environment was either non-significant or significant and in the same direction as plasticity.

#### Diffuse selection

To test whether warming changed patterns of diffuse selection, we compared selection gradients in plants pollinated by bumblebees and butterflies (generalized treatment; β_O_) with the gradients expected under additive selection (β_ADD_) (Sahli & Conner, 2011; Knauer & Schiestl, 2017). First, we estimated linear additive selection gradients for each trait by summing the gradients for plants pollinated by bumblebees or butterflies alone (terHorst *et al*., 2015). Then, we used β_ADD_ to estimate relative fitness (RF_ADD_) values under additive selection for each plant from the generalized treatment (see supplementary information). This resulted in a “predicted additive selection” dataset (“ADD”) with each plant from the generalized treatment having observed trait values and estimated relative fitness values. We merged the ADD dataset and the observed dataset from the generalized treatment. Finally, we fitted models (separate models for visual and scent traits, ambient and hot environment) with relative fitness as response variable, and floral traits, dataset (ADD or observed) and all possible interactions as fixed factors, and replicate as random factor (Knauer & Schiestl, 2017). A significant interaction between floral traits and dataset indicates nonadditive selection. We did not perform these analyses for nonlinear selection gradients because summing nonlinear selection gradients does not necessarily show additivity.

## Results

### Effect of temperature on plant traits

Elevated temperature had a significant effect on all measured phenological, visual and reward traits except for relative petal area of UV absorbance (Table S3). Hot-grown plants flowered 1-2 days earlier, were 3.4% taller (Fig. 1A), had 12.8% more leaves, and 21.4% more flowers (Fig. 1B) than ambient-grown plants. Flowers from hot-grown plants were smaller (Fig. 1C, reduced diameter (17.5%), petal area (35.8%), petal width (14.7%), and length (22.8%)) and produced 38.8% less nectar than flowers from ambient-grown plants. The emission of all quantified flower scent compounds was significantly affected by elevated temperature (Table S3). Hot-grown plants emitted less total floral volatiles and less of most individual floral scent compounds than ambient-grown plants (Fig. 1E-1F, Table S3). Nevertheless, hot-grown plants emitted 22.2% more benzaldehyde (Fig. 1D) and 143% more 1-butene-4-isothiocyanate than ambient-grown plants (Table S3).

**Figure 1.**
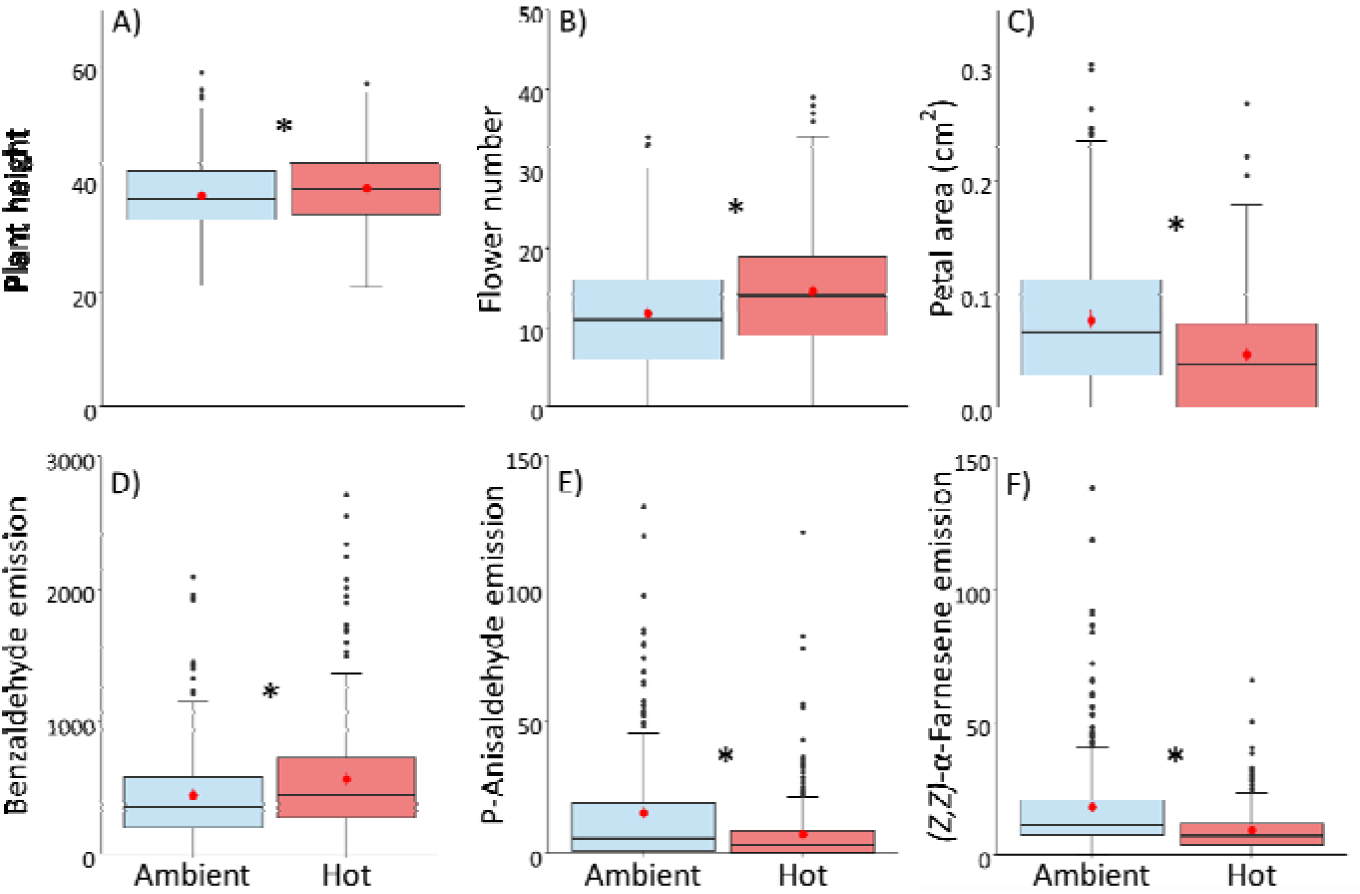
Effect of elevated temperature on A) plant height, B) number of flowers, C) petal area, D) benzaldehyde emission, E) p-anisaldehyde emission and E) *(Z,Z)-*α-farnesene emission of *Brassica rapa*. Scent emission is expressed as pg / flower / l / h. Plants were grown in an ambient (23°C, blue) or hot (27°C+ weekly 30°C day, red) temperature environment for 14 days prior to being measured. Boxplots show median (line), mean (red dot), first and third quartiles, and minimum and maximum. Outliers (1.5 times the interquartile range below the first quartile or above the third quartile) are represented by circles. Asterisks indicate a significant difference (*P* ≤ 0.05). Number of plant individuals per temperature varied between 281-293.

### Effect of temperature on insect behaviour

Temperature did not affect bumblebee flower visitation in the *Bombus* or generalized treatments (Table S4). In contrast, butterfly visitation was influenced by temperature in both *Pieris* and generalized treatments. For both treatments, hot-grown received more plant and flower visits, and more eggs compared to ambient-grown plants (Table S4). Temperature changed which plant traits influenced pollinator visitation and oviposition for each pollination treatment (Table S5,S6): Bumblebee and butterfly visitation and oviposition were influenced by different traits in the ambient and hot environment, which is detailed in the supplementary information (Table S7,S8).

### Effect of temperature on plant reproductive output

Temperature influenced plant reproduction for all pollination treatments (Table 1). For *Bombus*-treated plants, hot-grown plants produced less seeds per fruit compared to ambient-grown plants (Table 1). For *Pieris*-treated plants, hot-grown plants produced more seeds (15.5%) and fruits (14.0%), of which 339.5% more were damaged, but less seeds per fruit (5.9%) compared to ambient-grown plants (Table 1). Within the generalized treatment, hot-grown plants produced more seeds (36.8%) and fruits (27.2%), of which 264.6% more were damaged, compared to ambient-grown plants (Table 1).

**Table 1.** The effects of temperature and biotic treatment on plant reproduction. Mean reproductive traits and output of (generalized) linear (mixed) models showing effects of elevated temperature and three biotic treatments on *Brassica rapa* reproduction. Plants were exposed to elevated temperatures for 14 days after which pollination took place. *Bombus*-treated plants were visited by bumblebees, *Pieris*-treated plants were visited by *Pieris rapae* butterflies, and generalized-treated plants were visited by bumblebees and butterflies. Temperature treatments consisted of ambient (23°C) and hot (27°C+ weekly 30°C day). Bold values indicate results where *P* ≤ 0.05. Italic values indicate results where *P* ≤ 0.1.

|  |  |  | Ambient | Hot | Model |  |  |
| --- | --- | --- | --- | --- | --- | --- | --- |
| | | N | mean ± SE | mean ± SE | $\chi^2$ | df | P |
| *Bombus*-treated plants | fruit set | 79 | 0.75 ± 0.04 | 0.66 ± 0.05 | 1.8 | 1 | 0.184 |
| | fruits per plant | 81 | 13 ± 1.9 | 11 ± 1.3 | 2.5 | 1 | 0.117 |
| | seeds per plant | 81 | 129 ± 27 | 80 ± 14 | 2.1 | 1 | 0.149 |
| | seeds per fruit | 81 | 8.2 ± 0.9 | 6.2 ± 0.5 | 3.7 | 1 | 0.053 |
| *Pieris*-treated plants* | fruit set | 189 | 0.61 ± 0.03 | 0.70 ± 0.02 | 3.7 | 1 | 0.055 |
| | fruits per plant | 190 | 20. ± 1.2 | 22.8 ± 1.0 | 4.3 | 1 | **0.039** |
| | seeds per plant | 190 | 226 ± 16 | 261 ± 15 | 5.4 | 1 | **0.020** |
| | seeds per fruit | 190 | 11.9 ± 1.3 | 11.2 ± 0.5 | 6.9 | 1 | **0.008** |
| | damaged fruits (%) | 189 | 0.22 ± 0.05 | 0.64 ± 0.11 | 17.7 | 1 | **<0.001** |
| Generalized-treated plants | fruit set | 185 | 0.87 ± 0.02 | 0.91 ± 0.02 | 3.5 | 1 | 0.061 |
| | fruits per plant | 192 | 16 ± 1.1 | 20.67 ± 0.93 | 15.0 | 1 | **< 0.001** |
| | seeds per plant | 193 | 177 ± 14 | 242.32 ± 13.68 | 16.5 | 1 | **< 0.001** |
| | seeds per fruit | 192 | 11.1 ± 0.4 | 11.72 ± 0.41 | 1.2 | 1 | 0.269 |
| | damaged fruits | 193 | 0.03 ± 0.01 | 0.10 ± 0.02 | 26.5 | 1 | **< 0.001** |
|  | (%) |  |  |  |  |  |  |

\*Data from Rusman *et al.* (2025b).

### Temperature influences linear and non-linear selection

Linear selection gradients varied with temperature for all biotic treatments (Table S9). Plants visited by bumblebees in the ambient environment were under positive selection for flower number, and under negative selection for nectar per flower and phenylacetaldehyde emission (Fig. 2, Table S10). In the hot environment, plants were under positive selection for petal area and benzyl nitrile emission, and under negative selection for phenylacetaldehyde emission (Fig. 2, Table S10).

**Figure 2.**
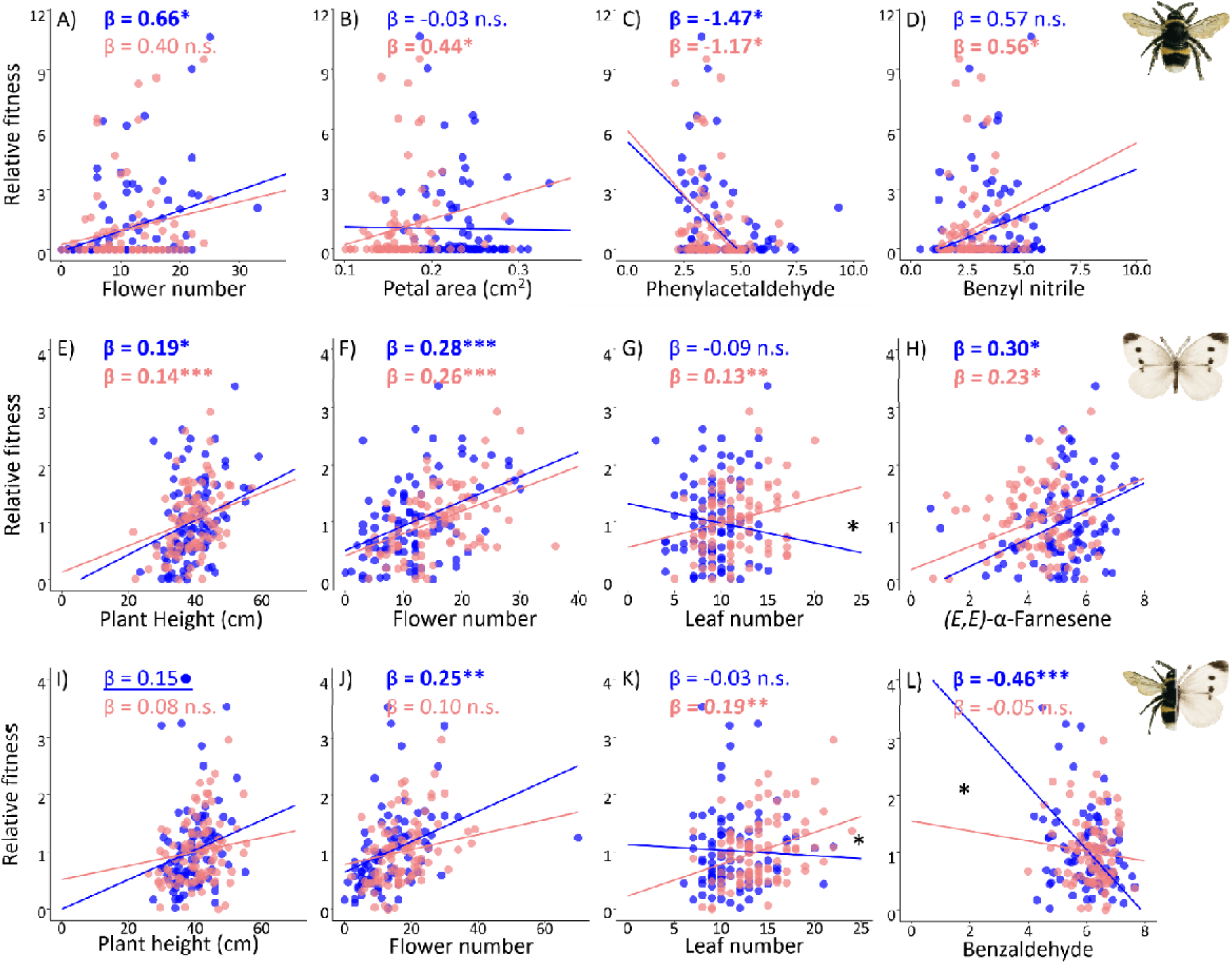
The effect of temperature on the directional selection of plant flowering traits. Scatterplots showing standardized selection gradients and regression lines for linear selection on plant visual traits of plants grown and tested in the ambient (23°C, blue) and hot (27°C+ weekly 30°C day, red) environment under different biotic treatments. **A-D**: Selection for *Bombus*-treated plants acting on A) flower number, B) petal area C) phenylacetaldehyde emission and D) benzyl nitrile emission. **E-H**: Selection for *Pieris*-treated plants acting on E) plant height, F) flower number G) leaf number and H) *(E,E)-*α-farnesene emission. **I-L**: Selection for generalized-treated plants acting on I) plant height, J) flower number, K) leaf number and L) benzaldehyde emission. Bold values indicate results where *P* ≤ 0.05 or the 95% C.I. does not include zero. Underlined values indicate results where *P* ≤ 0.1. Asterisks and dots indicate: *P* ≤ 0.05, with * P < 0.05, ** P < 0.01, *** P < 0.001, ● *P* ≤ 0.1. An asterisk between regression lines indicates an interactive effect of temperature on selection. Scent emission is ln (x + 1) transformed and expressed as pg / flower / l / h.

For plants visited by butterflies, there was an interactive effect of temperature on selection for leaf number: Leaf number was under positive directional selection in the hot environment but not in the ambient environment (Fig. 2, Table S9-S10). Plant height, flower number and *(E,E)-*α-farnesene emission were under positive selection in both environments (Fig. 2, Table S9,S10).

For plants visited by both insects, there was an interactive effect of temperature on selection for leaf number and benzaldehyde emission: leaf number was under positive selection in the hot but not the ambient environment, while benzaldehyde was under negative selection in the ambient but not the hot environment (Fig. 2, Table S9-S10). In the ambient environment, plants were under positive selection for flower number (Fig. 2, Table S10).

Nonlinear selection gradients and directional selection differentials differed between temperature environments for all biotic treatments (see supplementary results, Table S11,S12).

### Selection on temperature-mediated plasticity

Selection on temperature-mediated plasticity varied with biotic treatment and was measured as 1) selection on the reaction norm, or 2) the adaptive value of plasticity. For plants visited by bumblebees, we found positive selection on the plasticity of benzaldehyde emission (plastic increase) and phenylacetaldehyde (plastic decrease, Fig. 3A, Table S13). Temperature-mediated plasticity for petal area and benzyl nitrile emission was maladaptive (Table 2). For plants visited by butterflies, we found negative selection on the temperature-mediated plasticity of nectar per flower (plastic decrease) and stabilizing selection on the plasticity of benzaldehyde emission (plastic increase, Fig. 3B, Table S13,S14). Temperature-mediated plasticity for leaf number was adaptive (Table 2). For plants visited by both butterflies and bumblebees we found positive selection on the temperature-mediated plasticity of leaf number (plastic increase) and stabilizing selection on the plasticity of p-anisaldehyde emission (plastic decrease; Fig. 3C, Table S13,S14). Temperature-mediated plasticity for leaf number was adaptive (Table 2).

**Figure 3.**
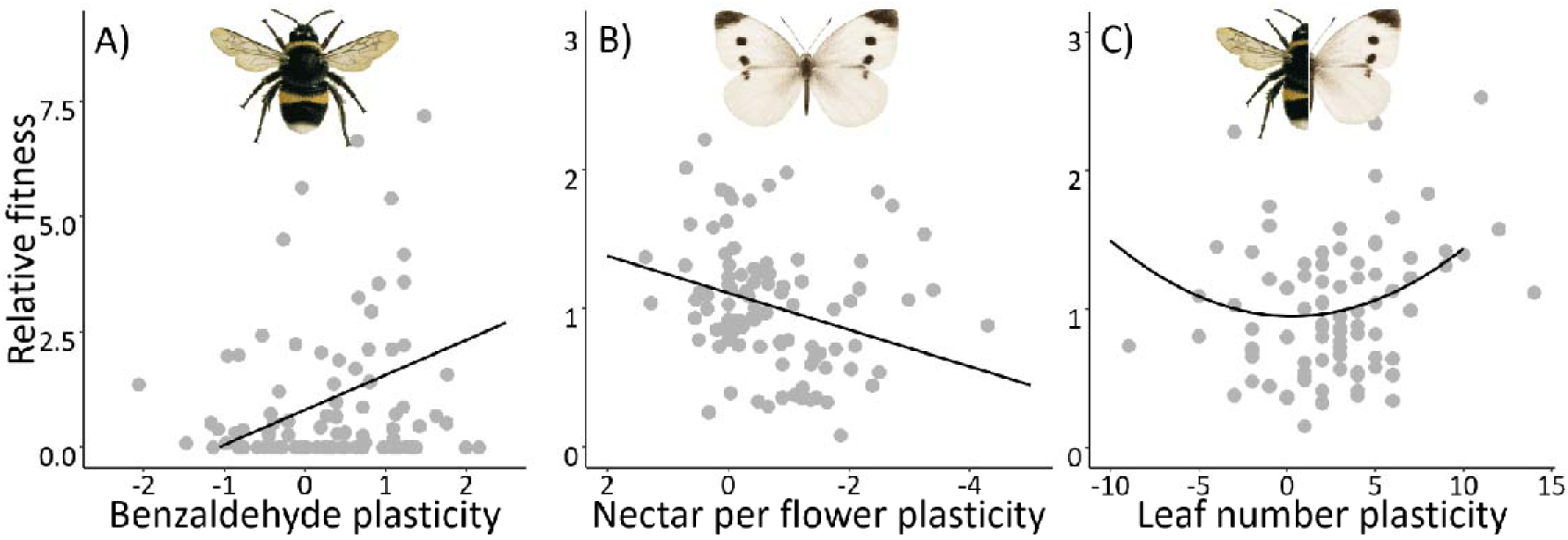
Selection on temperature-mediated plasticity. A: Selection on the temperature reaction norms of benzaldehyde emission in *Bombus*-treated plants. B: Selection on the temperature reaction norms of nectar per flower in *Pieris*-treated plants. C: Selection on the temperature reaction norms of leaves per plant in generalized-treated plants. Reaction norms were calculated comparing trait values of full-sibling plants grown in an ambient (23°C) and hot (27°C+ weekly 30°C day) temperature environment. In cases where the average reaction norm was negative (trait decreased with temperature) the X-axis was reversed for clarity.

**Table 2.** Adaptiveness of temperature-mediated plasticity for *Brassica rapa* plants. Direction of plasticity was positive (+) or negative (-) when the trait significantly increased or decreased in the hot (27°C+ weekly 30°C day) compared to the ambient (23°C) temperature environment. Treatment consisted of plant visitation by *Bombus terrestris* bumblebees*, Pieris rapae* butterflies, or both (generalized). Selection in the ambient temperature was always neutral (O), while selection in the hot environment was always positive (+), based on linear multiple regression analyses of visual or scent traits. We considered temperature-mediated plasticity adaptive (A) when selection in the hot environment was significant and in the same direction as plasticity, while selection in the ambient environment was either non-significant or significant and in the opposite direction as plasticity. We considered temperature-mediated plasticity maladaptive (M) when selection in the hot environment was significant and in the opposite direction of plasticity, while selection in the ambient environment was either non-significant or significant and in the same direction as plasticity. We only displayed traits that showed adaptive or maladaptive plasticity.

| Trait | Direction of plasticity | Treatment | Selection Ambient | Selection Hot | Adaptive? |
| --- | --- | --- | --- | --- | --- |
| Petal area (cm<sup>2</sup>) | - | Bombus | O | + | M |
| Benzyl nitrile emission | - | Bombus | O | + | M |
| Leaf number | + | Pieris | O | + | A |
| Leaf number | + | Generalized | O | + | A |

### Diffuse selection

Temperature changed diffuse linear selection for certain plant traits (Table S15). In the ambient but not in the hot environment, linear selection on four traits was nonadditive (Table S15): petal area (β_ADD_>β_O_: the selection gradient expect under additive selection was higher than the observed selection gradient), benzaldehyde emission (β_ADD_>β_O_), 2-amino benzaldehyde emission (β_ADD_<β_O_), and methyl anthranilate emission (β_ADD_>β_O_). In the hot environment but not the ambient environment, linear selection on four traits was nonadditive (Table S15): flowering time (β_ADD_>β_O_), UV surface area (β_ADD_>β_O_), 1-butene-4-isothiocyanate emission (β_ADD_>β_O_), and indole emission (β_ADD_<β_O_). In both environments, linear selection on nine traits was nonadditive: plant height, flower and leaf number, flower size, nectar per flower, and four floral scent compounds (Table S16).

Linear selection on the plasticity of 18 traits was nonadditive (Table S16): flowering time (β_ADD_ > β_O_), plant height (β_ADD_>β_O_), flower number (β_ADD_<β_O_ ), leaf number (β_ADD_<β_O_), flower size (β_ADD_<β_O_ ), petal area (β_ADD_>β_O_), UV surface area (β_ADD_<β_O_), and the emission of 11 floral scent compounds (Table S16).

## Discussion

Under global warming, increasing temperatures may impact plant evolutionary trajectories by altering natural selection (Jump & Peñuelas, 2005). However, how warming impacts selection and plant evolution may crucially depend on the composition of the surrounding biotic communities (Kühsel & Blüthgen, 2015; Nomoto *et al*., 2024). In our study, elevated temperature changed patterns of selection on plant traits and their plasticity, and the magnitude of these effects was highly dependent on community context. When multiple interactors were present, elevated temperature changed diffuse selection, showing that community context and warming interact to determine complex patterns of selection. In our follow-up studies, we found that these interactive effects of biotic interactors and temperature led to divergent (co)evolutionary trajectories (Rusman *et al*., 2025a; Traine *et al*., 2026a; Traine *et al*., 2026b). One of these studies investigated the evolutionary changes after 6 generations of selection under the same conditions as in the current study (Traine *et al*., 2026b). We found that the effects of warming on the evolution of plant traits and their plasticity depended heavily on community context, and that warming influenced nonadditive evolution in a multispecies community (Traine *et al*., 2026b). Hence, warming and community context influence complex selection patterns on plant flowering traits and their plasticity, with likely consequences for plant evolution.

Elucidating the contribution of abiotic and biotic factors to selective landscapes is important to understand evolutionary responses to environmental change. Biotic factors have long been considered as the main drivers of natural selection, but meta-analyses have shown that abiotic factors are as important for selection (Caruso *et al*., 2017; Caruso *et al*., 2019), even on a global scale (Siepielski *et al*., 2017). Interactive effects occur when abiotic and biotic factors impose selection on the same traits, or influence how traits respond to selection imposed by one another (Nomoto *et al*., 2024). Such interactive effects are likely common but are poorly understood. We show interactive effects of abiotic and biotic factors on selection: temperature altered pollinator-specific selection on plant flowering traits. When abiotic factors induce strong phenotypic plasticity, such plasticity can influence selection by changing trait-fitness relationships (Ehrlén & Valdés, 2020; Siepielski *et al*., 2022). Indeed, temperature induced strong phenotypic plasticity, and earlier work has shown that such plasticity can influence biotically-mediated selection (Gómez *et al*., 2024; Traine *et al*., 2024). This is not unique for temperature; soil-induced plasticity has been shown to influence pollinator-mediated selection as well (Dorey & Schiestl, 2022). Thus, interactive effects of abiotic and biotic factors can change patterns of phenotypic selection with plasticity potentially playing a prominent role.

Phenotypic plasticity itself can be under selection. Here, we show that selection on temperature-mediated plasticity differed between biotic interactions. For instance, plastic temperature-mediated increase in leaf number was under positive selection only for plants visited by both bumblebees and butterflies. Having more leaves may have helped tolerate increased herbivory under elevated temperatures (Kingsolver, 2000; Rusman *et al*., 2025b), and/or detrimental direct effects of temperature. Net photosynthetic rates in Brassicaceae can decrease above 25°C (Oh *et al*., 2015), so more leaves may compensate for reduced photosynthetic efficiency (Lawlor, 1995). Plasticity was not always under positive selection, however. Bumblebees preferred large flowers in the hot environment, leading to strong positive selection for this trait, while flower size plasticity was in opposite direction, *i.e.* heat induced smaller flowers, leading to floral plasticity in flower size being maladaptive. Reduction in flower size may present advantages in terms of thermoregulation at high temperatures (van der Kooi *et al*., 2019), but comes with costs in terms of reduced pollinator visitation (Descamps *et al*., 2021; Cordeiro & Dötterl, 2023; Traine *et al*., 2024). A similar effect was found for seasonal plasticity in *Moricandia arvensis*; summer morphs produced smaller flowers while pollinator-mediated selection favoured increased flower size (Gómez *et al*., 2024). Overall, our results suggest that temperature-mediated trait plasticity can evolve to be maintained/increased or be reduced depending on the biotic environment and potential ecological costs.

When environmental factors change plant interactions with multiple selection agents, changes in patterns of diffuse selection are predicted. Despite this, variation in diffuse selection has not been investigated in this context. Here, we show that elevated temperature changes patterns of diffuse selection on flower traits. Temperature can alter nonadditive selection by changing how butterflies influence selection imposed by bumblebees and vice versa. Butterflies and bumblebees can influence selection imposed by each other by direct ecological interactions, effects on plant fitness, and imposing selection on similar or correlated traits (Haloin & Strauss, 2008; Knauer & Schiestl, 2017). In our study, elevated temperature reduced plant fitness of bumblebee-pollinated plants while increasing the fitness of butterfly-pollinated plants. The increased contribution of butterflies to plant fitness for generalized-treated plants in the hot environment likely contributed to altered patterns of nonadditive selection. Furthermore, elevated temperatures changed the identity of the traits under selection by both pollinators and hence the opportunity for nonadditive selection to occur. Interestingly, we found nonadditive effects on several traits which were not under significant selection in the generalized treatment. This is not surprising, because in many cases additive selection predicted positive or negative selection on a trait, but this was diminished by nonadditive effects. Hence, nonadditive effects seem to reduce selection imposed by multiple selection agents (Afkhami *et al*., 2021), and such effects are further altered by environmental factors such as elevated temperature.

To gain deeper insights about the evolutionary consequences of climate change and species persistence, it is important to understand how natural selection is affected by phenomena such as warming within a community context (Jump *et al*., 2006; Anderson *et al*., 2025). In our study, we showed that increased temperatures changed selection on plant traits, that temperature-mediated trait plasticity was under varying selection, and that increased temperatures changed patterns of diffuse selection. Importantly, changes in selection varied widely depending on the biotic interactions plants experienced. Climate change is causing range shifts in both plants and insects, as well as facilitating species extinction and invasion events, so it is likely that plants will be exposed to rising temperatures and novel biotic communities (Rafferty, 2017; Anderson & Song, 2020). The interactive effects of novel biotic networks and temperature will ultimately shape patterns of phenotypic selection and evolutionary trajectories that lead to climate adaptation or extinction.

## Acknowledgements

We thank Franz Huber for helping with the analysis and quantification of flower scent, Laura Dallenbach for rearing the insects and helping with various tasks, Maeve Lin and Kajsa Bornhauser for helping with butterfly behavioural observations, Julia Weidmann for her help with counting fruits and seeds, Rayko Jonas and Markus Meierhofer for help with growing and caring for the plants. This research was funded by the Swiss National Science Foundation to FPS (grant No. 310030_208158).

## Author contributions

JT, FPS, and QR designed the study. JT and QR performed the experiments. JT and QR analysed the data. JT, FPS, and QR wrote the paper.

## Data availability

The raw data and code are available upon acceptance of the manuscript.

## Supporting information

Additional supporting information may be found online in the Supporting Information section at the end of the article.

**Methods S1.** Detailed description of insect rearing and origin, measurements of plant flowering phenology, morphology, reward, and scent, and statistical analyses.

**Results S1.** Detailed description of results related to flower traits and insect attraction, nonlinear selection gradients, and selection differentials.

**Methods S2.** Description of methods of experiment 2: Swapping temperature treatments between greenhouse cabins.

**Results S2.** Description of results of experiment 2: Swapping temperature treatments between greenhouse cabins.

**Table S1.** Results of experiments 1 and 2 showing plastic changes of *Brassica rapa* traits, insect behaviour, and plant reproduction in response to temperature.

**Table S2.** Results of experiment 1 and 2 showing the effect of temperature on linear selection differentials.

**Table S3.** Plastic change of *Brassica rapa* traits in response to temperature.

**Table S4.** Changes in bumblebee and butterfly behaviour in response to temperature.

**Table S5.** Interactive effects of temperature and plant traits on pollinator visitation.

**Table S6.** Interactive effects of temperature and plant traits on oviposition.

**Table S7.** Effects of plant traits on pollinator visitation in different temperature environments.

**Table S8.** Effects of plant traits on oviposition in different temperature environments.

**Table S9.** The interactive effects of temperature and biotic treatments on the directional selection of plant traits.

**Table S10.** The effects of temperature on the directional selection of plant traits under different biotic treatments.

**Table S11.** The effects of temperature on the quadratic selection of plant traits under different biotic treatments.

**Table S12.** The effects of temperature on the selection differentials of plant traits under different biotic treatments.

**Table S13.** The directional selection of temperature-mediated plasticity under different biotic treatments.

**Table S14.** The non-linear selection of temperature-mediated plasticity under different biotic treatments.

**Table S15.** Effects of temperature on patterns of additive and nonadditive linear selection.

**Table S16.** Effects of temperature on patterns of additive and nonadditive selection on temperature-mediated plasticity.

## References

Afkhami ME, Friesen ML, Stinchcombe JR. 2021. Multiple mutualism effects generate synergistic selection and strengthen fitness alignment in the interaction between legumes, rhizobia and mycorrhizal fungi. Ecology Letters 24: 1824–1834.

Agren J, Schemske DW. 1993. The cost of defense against herbivores: An experimental study of trichome production in *Brassica rapa*. The American Naturalist 141: 338–350.

Anderson JT, DeMarche ML, Denney DA, Breckheimer I, Santangelo J, Wadgymar SM. 2025. Adaptation and gene flow are insufficient to rescue a montane plant under climate change. Science 388: 525–531.

Anderson JT, Song B-H. 2020. Plant adaptation to climate change—Where are we? Journal of Systematics and Evolution 58: 533–545.

Atmowidi TRI, Buchori D, Manuwoto S, Suryobroto B, Hidayat P. 2007. Diversity of pollinator insects in relation to seed set of mustard (*Brassica rapa* L.: Cruciferae). HAYATI Journal of Biosciences 14: 155–161.

Becklin KM, Anderson JT, Gerhart LM, Wadgymar SM, Wessinger CA, Ward JK. 2016. Examining plant physiological responses to climate change through an evolutionary lens Plant Physiology 172: 635–649.

Borghi M, Perez de Souza L, Yoshida T, Fernie AR. 2019. Flowers and climate change: a metabolic perspective. New Phytologist 224: 1425–1441.

Campbell DR, Powers JM. 2015. Natural selection on floral morphology can be influenced by climate. Proceedings of the Royal Society B: Biological Sciences 282: 20150178.

Caruso CM, Eisen KE, Martin RA, Sletvold N. 2019. A meta-analysis of the agents of selection on floral traits. Evolution 73: 4–14.

Caruso CM, Martin RA, Sletvold N, Morrissey MB, Wade MJ, Augustine KE, Carlson SM, MacColl ADC, Siepielski AM, Kingsolver JG. 2017. What are the environmental determinants of phenotypic selection? A meta-analysis of experimental studies. The American Naturalist 190: 363–376.

Cordeiro GD, Dötterl S. 2023. Global warming impairs the olfactory floral signaling in strawberry. BMC Plant Biology 23: 549.

de Manincor N, Fisogni A, Rafferty NE. 2023. Warming of experimental plant–pollinator communities advances phenologies, alters traits, reduces interactions and depresses reproduction. Ecology Letters 26: 323–334.

Descamps C, Jambrek A, Quinet M, Jacquemart A-L. 2021. Warm temperatures reduce flower attractiveness and bumblebee foraging. Insects 12: 493.

Dorey T, Schiestl FP. 2022. Plant phenotypic plasticity changes pollinator-mediated selection. Evolution 76: 2930–2944.

Ehrlén J, Valdés A. 2020. Climate drives among-year variation in natural selection on flowering time. Ecology Letters 23: 653–662.

Fischer AM, Strassmann KM, Croci-Maspoli M, Hama AM, Knutti R, Kotlarski S, Schär C, Schnadt Poberaj C, Ban N, Bavay M, et al. 2022. Climate scenarios for Switzerland CH2018 – Approach and implications. Climate Services 26: 100288.

Frane AV. 2021. Experiment-wise type I error control: A focus on 2 × 2 designs. Advances in Methods and Practices in Psychological Science 4: 10.1177/2515245920985137.

García-Pérez MA. 2023. Use and misuse of corrections for multiple testing. Methods in Psychology 8: 100120.

Gérard M, Vanderplanck M, Wood T, Michez D. 2020. Global warming and plant–pollinator mismatches. Emerging Topics in Life Sciences 4: 77–86.

Gómez JM, González-Megías A, Armas C, Narbona E, Navarro L, Perfectti F. 2024. Selection maintains a nonadaptive floral polyphenism. Evolution Letters 8: 610–621.

Gómez JM, Navarro L, González-Megías A, Armas C, Perfectti F, Caravantes Á, Sánchez R. 2025. Pollination effectiveness affects the level of generalisation of a plant species with phenotypically plastic flowers. AoB PLANTS 17.

Gómez JM, Perfectti F, Armas C, Narbona E, González-Megías A, Navarro L, DeSoto L, Torices R. 2020. Within-individual phenotypic plasticity in flowers fosters pollination niche shift. Nature Communications 11: 4019.

Haloin JR, Strauss SY. 2008. Interplay between ecological communities and evolution. Annals of the New York Academy of Sciences 1133: 87–125.

Hamann E, Blevins C, Franks SJ, Jameel MI, Anderson JT. 2021. Climate change alters plant–herbivore interactions. New Phytologist 229: 1894–1910.

Hunter DC, Pemberton JM, Pilkington JG, Morrissey MB. 2018. Quantification and decomposition of environment-selection relationships. Evolution 72: 851–866.

Janzen DH. 1980. When is it coevolution? Evolution 34: 611–612.

Jump AS, Hunt JM, MartÍNez-Izquierdo JA, PeÑUelas J. 2006. Natural selection and climate change: temperature-linked spatial and temporal trends in gene frequency in *Fagus sylvatica*. Molecular Ecology 15: 3469–3480.

Jump AS, Peñuelas J. 2005. Running to stand still: adaptation and the response of plants to rapid climate change. Ecology Letters 8: 1010–1020.

Kingsolver JG. 2000. Feeding, Growth, and the Thermal Environment of Cabbage White Caterpillars, *Pieris rapae* L. Physiological and Biochemical Zoology 73: 621–628.

Kingsolver JG, Pfennig DW. 2007. Patterns and power of phenotypic selection in nature. Bioscience 57: 561–572.

Knauer AC, Schiestl FP. 2017. The effect of pollinators and herbivores on selection for floral signals: a case study in *Brassica rapa*. Evolutionary Ecology 31: 285–304.

Kühsel S, Blüthgen N. 2015. High diversity stabilizes the thermal resilience of pollinator communities in intensively managed grasslands. Nature Communications 6: 7989.

Lande R, Arnold SJ. 1983. The measurement of selection on correlated characters. Evolution 37: 1210–1226.

Lawlor DW. 1995. Photosynthesis, productivity and environment. Journal of Experimental Botany 46: 1449–1461.

Morrissey MB, Ruxton GD. 2018. Multiple regression is not multiple regressions: The meaning of multiple regression and the non-problem of collinearity. *Philosophy*, Theory, and Practice in Biology 10.

Napier JD, Heckman RW, Juenger TE. 2023. Gene-by-environment interactions in plants: Molecular mechanisms, environmental drivers, and adaptive plasticity. The Plant Cell 35: 109–124.

Nomoto H, Fior S, Alexander J. 2024. Competitors alter selection on alpine plants exposed to experimental climate change. Evolution Letters 8: 114–127.

Oh S, Moon KH, Song EY, Son I-C, Koh SC. 2015. Photosynthesis of Chinese cabbage and radish in response to rising leaf temperature during spring. *Horticulture*, Environment, and Biotechnology 56: 159–166.

Perneger TV. 1998. What’s wrong with Bonferroni adjustments. BMJ 316: 1236.

Rader R, Howlett BG, Cunningham SA, Westcott DA, Newstrom-Lloyd LE, Walker MK, Teulon DA, Edwards W. 2009. Alternative pollinator taxa are equally efficient but not as effective as the honeybee in a mass flowering crop. Journal of Applied Ecology 46: 1080–1087.

Rafferty NE. 2017. Effects of global change on insect pollinators: multiple drivers lead to novel communities. Current Opinion in Insect Science 23: 22–27.

Ramos SE, Schiestl FP. 2020. Herbivory and pollination impact on the evolution of herbivore-induced plasticity in defense and floral traits. Evolution Letters 4: 556–569.

Rubin M. 2021. When to adjust alpha during multiple testing: a consideration of disjunction, conjunction, and individual testing. Synthese 199: 10969–11000.

Rusman Q, Figueira T, Traine J, Schiestl FP. 2025a. Coevolution in a warming world: an experimental test of the geographic mosaic of coevolution. bioRxiv: 686912.

Rusman Q, Traine J, Schiestl FP. 2025b. Elevated temperature diminishes reciprocal selection in an experimental plant-pollinator-herbivore system. Ecology Letters 28: e70060.

Sahli HF, Conner JK. 2011. Testing for conflicting and nonadditive selection: Floral adaptation to multiple pollinators through male and female fitness. Evolution 65: 1457–1473.

Scaven VL, Rafferty NE. 2013. Physiological effects of climate warming on flowering plants and insect pollinators and potential consequences for their interactions. Current Zoology 59: 418–426.

Siepielski AM, Gómez-Llano M, McPeek MA. 2022. Environmental conditions during development affect sexual selection through trait-fitness relationships. The American Naturalist 199: 34–50.

Siepielski AM, Morrissey MB, Buoro M, Carlson SM, Caruso CM, Clegg SM, Coulson T, DiBattista J, Gotanda KM, Francis CD, et al. 2017. Precipitation drives global variation in natural selection. Science 355: 959–962.

Strauss SY, Irwin RE. 2004. Ecological and evolutionary consequences of multispecies plant-animal interactions. Annual Review of Ecology, Evolution, and Systematics 35: 435–466.

Strauss SY, Sahli H, Conner JK. 2005. Toward a more trait-centered approach to diffuse (co)evolution. New Phytologist 165: 81–90.

Sztepanacz JL, Houle D. 2024. Regularized regression can improve estimates of multivariate selection in the face of multicollinearity and limited data. Evolution Letters 8: 361–373.

terHorst CP, Lau JA, Cooper IA, Keller KR, Rosa RJL, Royer AM, Schultheis EH, Suwa T, Conner JK. 2015. Quantifying nonadditive selection caused by indirect ecological effects. Ecology 96: 2360–2369.

terHorst CP, Zee PC, Heath KD, Miller TE, Pastore AI, Patel S, Schreiber SJ, Wade MJ, Walsh MR. 2018. Evolution in a community context: Trait responses to multiple species interactions. The American Naturalist 191: 368–380.

Toftegaard T, Posledovich D, Navarro-Cano JA, Wiklund C, Gotthard K, Ehrlén J. 2019. Butterfly–host plant synchrony determines patterns of host use across years and regions. Oikos 128: 493–502.

Traine J, Rusman Q, Schiestl FP. 2024. Too hot to handle: temperature-induced plasticity influences pollinator behaviour and plant fitness. New Phytologist 243: 1571–1585.

Traine J, Rusman Q, Schiestl FP. 2026a. Adaptive evolution can mitigate the negative effects of temperature stress on plant–pollinator interactions. New Phytologist 249: 554–568.

Traine J, Schiestl FP, Rusman Q. 2026b. Warming and community context drive rapid plant evolution. Functional Ecology 0: 1–15.

van der Kooi CJ, Kevan PG, Koski MH. 2019. The thermal ecology of flowers. Annals of Botany 124: 343–353.

Van Kleunen M, Fischer M. 2005. Constraints on the evolution of adaptive phenotypic plasticity in plants. New Phytologist 166: 49–60.

Wang S, Feng W-W, Liu M-C, Huang K, Arnold PA, Nicotra AB, Feng Y-L. 2022. Inherent conflicts between reaction norm slope and plasticity indices when comparing plasticity: a conceptual framework and empirical test. Oecologia 198: 593–603.

Zhang S-S, Yang H, Ding L, Song Z-T, Ma H, Chang F, Liu J-X. 2017. Tissue-specific transcriptomics reveals an important role of the unfolded protein response in maintaining fertility upon heat stress in *Arabidopsis*. The Plant Cell 29: 1007–1023.

Zhu H, Zou X, Wang D, Wan S, Wang L, Guo J. 2015. Responses of community-level plant-insect interactions to climate warming in a meadow steppe. Scientific Reports 5: 18654.

