## Supplemental information for "Warming and biotic interactions shape selection on plant traits and their plasticity"

### Methods S1

*Insect rearing and origin*

We kept adult butterflies and caterpillars in a greenhouse compartment (22±1°C, 60%RH, 16h:8h light:dark). Adults were housed in rearing cages (90 x 60 x 60 cm) and supplied with flowering fast-cycling *B. rapa* plants for feeding and oviposition, as well as supplementary water and nectar. First- and second-instar caterpillars were kept on fast-cycling *B. rapa*, but from their third instar we transferred them to Savoy cabbage leaves (*Brassica olaraceae* var. *sabauda*) where they remained until pupation. Information on the origin of the rearing can be found in Traine et al. (2024); Rusman et al. (2025).

We purchased two bumblebee hives (Hummelvolk Bombus Maxi, Andermatt Biocontrol, Switzerland) and kept them in flight cages (3 x 1 x 1 m) inside our greenhouse (22±1°C, 60%RH, 16h:8h light:dark). We supplied bumblebees flowering fast-cycling *Brassica rapa* plants, and supplementary nectar (BioGluc, Biobest, Belgium) and pollen (Blütenpollen Multiflora, KoRo Handels GmbH, Germany). To encourage foraging, feeding plants were removed two days prior to bioassays and supplementary nectar and pollen sources 16hs before(Dorey & Schiestl, 2024; Traine et al., 2024).

*Plant flowering phenology, morphology and rewards*

Two weeks after sowing, we began monitoring plants daily for the start of flowering. Four weeks after sowing (21-25 days), we measured plant height (in cm, with a 5 mm precision), number of open flowers and number of leaves. For flower size we took three flowers per plant and measured their diameter with a digital calliper. We then separated their petals from the pedicel and glued them with transparent scotch tape to an A4 paper sheet. We scanned these paper sheets at 600 DPI using a printer (Canon imageRunner Advance DX) and produced .JPEG images. Using the software ImageJ we used the scanned imaged to calculate petal area (in cm2), petal length (in cm) and petal width (in cm). The paper sheets with glued petals were also photographed under UV light (1W-UV Torch, λ= 365 nm and intensity = 6.6 mW/cm2) using a photo camera with an UV-transmission filter (Nikon D200 digital SLR, with quartz lens: Nikon UV Nikkor f4.5/105mm with Baader 2" U-Filter (310 - 390 nm UV transmission). We analysed the ensuing photographs on ImageJ to calculate the area of UV absorbance per petal (in $cm^2$), which was then divided by the petal area to calculate the relative area of UV absorbance per petal. We measured nectar volume from the same three flowers used for morphology. To extract nectar, we pressed a 5µl glass capillary against the nectaries. We measured the length of the capillary filled with nectar in mm using a digital calliper and then calculated the nectar volume in µl by comparing the measured capillary length with the length of the 1 µl mark in the capillary. A single capillary was used per three flowers (per plant), so the measurement was divided by three to calculate the nectar volume per flower.

*VOC collection and quantification*

We collected and quantified flower scent compounds four weeks after sowing (21-24 days) in the ambient temperature greenhouse cabin. We used headspace sorption with a push-pull system (Traine et al., 2024; Rusman et al., 2025). The main inflorescence of each plant was enclosed in a glass cylinder with a silanized surface (Sigmacote; Sigma-Aldrich, Buchs, Switzerland). The base of the glass cylinder was sealed using two Teflon plates, allowing only the inflorescence stem to pass through a 0.5 cm diameter hole. Air was introduced into the cylinder through a side opening, which was equipped with an activated charcoal filter to purify incoming air. Another side opening contained a glass tube filled with an absorbent (30 mg of Tenax), through which the air was pulled. Air was pushed and pulled by pumps at a rate of 120 ml/min. At the beginning of each collection, we recorded the number of flowers inside the glass cylinders. Around 25 plants were sampled per round, plus an empty glass cylinder used as air control. Collections lasted two hours and were performed in the morning or afternoon; plants were sampled only once. We quantified flower scent using chromatography with mass selective detection (GC–MSD).

Samples were injected into an Agilent 6890N GC system (Agilent Technologies, Santa Clara, CA, USA) via a MultiPurpose Sampler (MPS; Gerstel, Mülheim, Germany) possessing a thermal desorption unit (TDU) and a cold injection system (CIS). The GC was equipped with an HP-5 column (15 m length, 0.25 mm ID, 0.25 µm film thickness), through which helium (the carrier gas) could flow at a rate of 2 ml/min. Compound desorption from the Tenax happened by heating the TDU from 30 to 240°C at a rate of 60°C per minute, with a 5-minute hold at the final temperature. Then, rapid cooling to -150°C trapped the desorbed compounds in the CIS. For injection, the CIS was heated at a rate of 12°C per second until reaching 250°C and then held at that temperature for 3 minutes. Each compound was identified and quantified using a mass selective detector (Agilent MSD 5975). The mass spectra were compared to the National Institute of Standards and Technology (NIST) library and to the spectra of synthetic standards. Quantification of each compound, based on total ion counts, was done employing a calibration curve for target ions specific to each individual compound. All compounds were standardized to picograms per flower per liter of sampled air per hour.

*Statistical analyses*

*Insect attraction and plant traits*

To determine whether insects were attracted by different plant traits between temperature treatments, we performed generalized linear mixed models. For each biotic treatment (including *Pieris*-treated plants since these analyses were not performed in Rusman et al. (2025)) and plant trait separately, we tested first the interactive effects of temperature and each individual plant trait on number of flower visits or oviposition. Replicate and plant family were included as random factors. Second, we tested within each temperature the effect of each individual plant trait on number of flower visits or oviposition. Replicate was included as a random factor.

*Nonlinear selection analyses*

We obtained nonlinear selection gradients from models containing linear and quadratic terms (Palacio et al., 2019). The quadratic term coefficients were doubled to estimate stabilizing/disruptive selection gradients (Stinchcombe et al., 2008). Since the models included a high number of traits, we did not calculate correlational selection to avoid model overfitting (Chapurlat et al., 2019).

*Diffuse selection*

To test whether warming changed patterns of diffuse selection, we compared selection gradients in plants pollinated by bumblebees and butterflies (generalized treatment; $\beta_O$) with the gradients expected under additive selection ($\beta_{ADD}$) (Sahli & Conner, 2011; Knauer & Schiestl, 2017). First, we estimated linear additive selection gradients for each trait by summing the gradients for plants pollinated by bumblebees or butterflies alone (terHorst et al., 2015). Then, we used $\beta_{ADD}$ to estimate relative fitness ($RF_{ADD}$) values under additive selection for each plant from the generalized treatment. Relative fitness under additive selection was estimated from multivariate models (separate models for visual and scent traits) with $\beta_{ADD}$ for each trait, and the plant trait values from generalized-treated plants. Hence we used $RF_{ADD} = (\beta_{ADD1}*T_1) + (\beta_{ADD2}*T_2) + .... + (\beta_{ADDX}*T_X)$, where T is the trait value observed for generalized-treated plants, and x the total number of traits. This resulted in a "predicted additive selection" dataset ("ADD") with each plant from the generalized treatment having observed trait values and estimated relative fitness values. We merged the ADD dataset and the observed dataset from the generalized treatment and labelled each dataset accordingly (ADD or observed). Finally, we fitted models (separate models for visual and scent traits, ambient and hot environment) with relative fitness as response variable, and floral traits, dataset (ADD or observed) and all possible interactions as fixed factors, and replicate as random factor (Knauer & Schiestl, 2017). A significant interaction between floral traits and dataset indicates nonadditive selection. We did not perform these analyses for nonlinear selection gradients because summing nonlinear selection gradients does not necessarily show additivity.

*R-packages used*

We checked each model's distribution fit using the function *SimulateResiduals* from package "DARHMa" (Hartig, 2022), except for Cox models, where the proportional hazards assumption was checked using the function *cox.zph* from package "survival" (Therneau & Lumley, 2015). To perform (G)LMMs, we used the functions *lmer* and *glmer* from "lme4" (Bates et al., 2015), and *glmmTMB* from glmmTMB (Brooks et al., 2017). We used F-tests to derive p-values from LMMs and Wald chi-square tests to derive p-values from GLMMs. We obtained p-values using the *Anova* function from "car" (Fox & Weisberg, 2019). All figures were created using the function *ggplot* from the package "ggplot2" (Wickham, 2011). We calculated variance inflation factors (VIFs) using the *vif* function from package "car" (Fox & Weisberg, 2019). Bootstrapping was performed with the functions *boot* and *boot.ci* from package "boot" (Canty, 2002; Palacio et al., 2019).

**Results S1**

*Flower traits and insect attraction*

Temperature changed which flower traits influenced pollinator visitation and oviposition for each pollination treatment (Table S5,S6). For bumblebee visitation, temperature had an interactive effect with nectar per flower (Table S5). In the ambient environment, plant height, flower number, leaf number, and the emission of 2-amino benzaldehyde, benzyl nitrile, *(Z,Z)*-α-farnesene and *(E,E)*-α-farnesene influenced bumblebee visitation (Table S7). In the hot environment, plant height, flower number, flower size, and the emission of 3-hexenyl acetate influenced bumblebee visitation (Table S7).

For butterfly visitation in the *Pieris* treatment, temperature had an interactive effect with plant height and UV surface area (Table S5). Indeed, plant height and UV surface area influenced butterfly visitation in the ambient but not the hot environment (Table S7). In addition, in the ambient environment, flower number, flower size (flower diameter, petal area, length and width), and *(E,E)*-α-farnesene emission influenced butterfly visitation (Table S7). In the hot environment, flower number and flower size (petal area and width) influenced butterfly visitation (Table S7). For oviposition, temperature had an interactive effect with plant height, leaf number, and *(E,E)*-α-farnesene emission (Table S6). Indeed, plant height and leaf number influenced oviposition in the ambient but not hot environment (Table S8). In addition, in the ambient environment, *(Z)*-3-hexen-1-ol-acetate emission influenced oviposition (Table S8). In the hot environment, no traits correlated with oviposition (Table S8).

For pollinator visitation in the generalized treatment, temperature had an interactive effect with nectar per flower and *(Z)*-3-hexen-1-ol-acetate emission (Table S5). Indeed, nectar per flower influenced insect visitation in the hot environment only, while *(Z)*-3-hexen-1-ol-acetate emission influenced insect visitation in the ambient environment only (Table S7). In addition in the hot environment, petal area, width, and UV surface area influenced insect visitation (Table S7). In both environments, plant height, flower number, flower size (flower diameter), and the emission of benzaldehyde and 1-butene-4-isothiocyanate influenced insect visitation (Table S7). For oviposition, temperature had an interactive effect on the emission of methyl benzoate, p-anisaldehyde, methyl anthranilate, *(Z)*-3-hexen-1-ol-acetate, and *(Z,Z)*-α-farnesene (Table S6). Indeed, p-anisaldehyde and methyl anthranilate influenced oviposition in the ambient environment, while *(Z,Z)*-α-farnesene emission influenced oviposition in the hot environment (Table S8). In addition, in the ambient environment, plant height, petal area and width, indole and total volatile emission influenced oviposition (Table S8). In the hot environment, 1-butene-4-isothiocyanate emission influenced oviposition (Table S8).

*Nonlinear selection gradients*

Nonlinear selection gradients varied with temperature for all biotic treatments. For *Bombus*-treated plants in the ambient environment, plants were under stabilizing selection for flowering time, nectar per flower, and 3-hexenyl acetate emission, and under disruptive selection for leaf number, and the emission of (*Z,Z*)-α-farnesene (Table S11). In the hot environment, plants were under stabilizing selection for nectar per flower (Table S11). For *Pieris*-treated plants in the ambient environment, plants were under disruptive selection for leaf number (Table S11). In the hot environment, plants were under stabilizing selection for 1-butene-4-isothiocyanate emission (Table S11). For generalized-treated plants in the ambient environment, plants were under stabilizing selection for flower number, and under disruptive selection for UV surface area cover of petals (Table S11). In the hot environment, plants were under disruptive selection for benzyl nitrile emission (Table S11).

*Selection differentials*

Directional selection differentials differed between temperature environments for all pollination treatments. For *Bombus* treatments in the ambient environment, we detected positive selection differentials for plant height and flower number, and negative selection differentials for nectar per flower and phenylacetaldehyde emission (Table S12). In the hot environment we detected a positive selection differential for plant height, and negative selection differentials for phenylacetaldehyde emission (Table S12). For *Pieris* treatments in the ambient environment, we detected positive selection differentials for plant height, flower number, β-pinene and *(E,E)*-α-farnesene emission (Table S12). In the hot environment we detected positive selection differentials for plant height, flower number, leaf number, petal area and petal width (Table S12). For the generalized treatment in the ambient environment, we detected positive selection differentials for plant height and flower number, and negative selection differentials for benzaldehyde emission (Table S12). In the hot environment, we detected positive selection differentials for plant height, flower and leaf number, and flower diameter (Table S12).

**Experiment 2: Swapping temperature treatments between greenhouse cabins**

**Methods S2**

To show that greenhouse cabin and temperature treatment were independent, we swapped the temperature treatments between the greenhouse cabins. During the main experiment (referred to as "experiment 1"), greenhouse cabin 1 contained the ambient temperature treatment, while greenhouse cabin 2 contained the hot temperature treatment. For experiment 2, cabin 1 contained the hot temperature treatment, and greenhouse cabin 2 contained the ambient temperature treatment. We took seeds from the same full-sib families as used in experiment 1 and grew 294 plants. 147 plants were grown in the ambient greenhouse (2 sets of 49 plants of plant families 1-49, and one set of 49 plants of plant families 50-98), and 147 plants were grown in the hot greenhouse (2 sets of 49 plants of plant families 1-49, and one set of 49 plants of plant families 50-98). In each greenhouse cabin, we phenotyped 98 plants (1 plants of each of the 98 families). We measured plant height, leaf and flower number, flower diameter, petal area, length and width, and nectar amount 21-25 days after sowing, as described for experiment 1. These plants were subsequently used for bioassays with bumblebees and assessment of phenotypic selection. The remaining plants (one plant of families 1-49) were used in butterfly behaviour assays.

Bumblebee assays were performed as described for experiment 1; except we did not record bumblebee visitation time. Butterfly assays were performed as described for experiment 1, except all 25 butterflies were released simultaneously and thus plant visitations lasted a total of 2 days. Oviposition was not recorded, and plants were discarded at the end of the second day of plant and flower visitations. One month after the bioassays bumblebee-visited plants were moved to a different greenhouse for drying and fruit collection and plant reproduction was assessed as described for experiment 1.

*Statistical analysis*

To investigate the effect of temperature treatment and greenhouse cabin on plant flowering traits, insect behaviour, plant reproduction, and phenotypic selection, we merged data from experiment 1 with experiment 2 for the relevant variables (Table S1-S2). Similar for experiment 1, we run (generalized) linear mixed models. We included temperature, experiment, and their interaction as fixed factors, and replicate as random factor. Significance of temperature indicates that the temperature treatments affected the respective variable independent of the greenhouse cabins. Significance of experiment indicates that there were differences in the response variable between experiments. The most likely explanation is that the response variable differed between experiments due to external factors; performing greenhouse experiments in spring (experiment 2) or summer (experiment 1) will slightly influence plant growth and floral trait expression. A significant interaction between temperature and experiment indicates that the temperature treatment varied between the greenhouse cabins. Finally, we calculated selection differentials for experiment 2 similar as described for experiment 1, and compared the differentials found in both experiments.

### Results S2

Temperature influenced the same flowering traits in experiment 2 as in experiment 1, indicating that our temperature treatment was independent of the greenhouse cabins (Table S1). The same was true for insect behaviour: in experiment 2 bumblebee behaviour was not influenced by temperature, but butterfly behaviour was (Table S1). This indicates that both plants and insects responded similar to the temperature treatment independent of the cabins. Moreover, we found similar traits under selection in experiment 2 as in experiment 1 (Table S2), indicating that effects of our temperature treatment on selection were also independent of the greenhouse cabins.

For none of the plant flowering traits, insect behaviour, or plant reproduction, we found an interaction between experiment and temperature, indicating that our temperature treatment did not vary between the greenhouse cabins, except for nectar amount (Table S1). This was not because the temperature effects on nectar differed between the cabins, in both experiments elevated temperature reduced nectar amount (Table S1), but because the effect was stronger in experiment 2 compared to experiment 1.

Experiment influenced various plant flowering traits and insect behaviour (Table S1). This was most likely because the timing of the experiments (as in time of year) influenced plant growth: plants were smaller with less leaves and flowers in both cabins in experiment 2 (performed between February – March 2026) compared to the main experiment (performed between July – September 2021). This subsequently influenced insect visitation.

### References


Bates, D., Maechler, M., Bolker, B., & Walker, S. (2015). Fitting linear mixed-effects models using lme4. *Journal of Statistical Software, 67*, 1-48. doi:10.18637/jss.v067.i01

Brooks, M. E., Kristensen, K., Van Benthem, K. J., Magnusson, A., Berg, C. W., Nielsen, A., . . . Bolker, B. M. (2017). glmmTMB balances speed and flexibility among packages for zero-inflated generalized linear mixed modeling. *The R journal, 9*(2), 378-400.

Canty, A. J. (2002). Resampling methods in R: the boot package. *The Newsletter of the R Project Volume, 2*(3), 2-7.

Chapurlat, E., Ågren, J., Anderson, J., Friberg, M., & Sletvold, N. (2019). Conflicting selection on floral scent emission in the orchid *Gymnadenia conopsea*. *New Phytologist, 222*(4), 2009-2022. doi:10.1111/nph.15747

Dorey, T., & Schiestl, F. P. (2024). Bee-pollination promotes rapid divergent evolution in plants growing in different soils. *Nature Communications, 15*(1), 2703. doi:10.1038/s41467-024-46841-4

Fox, J., & Weisberg, S. (2019). *An R companion to applied regression*: Sage publications.

Hartig, F. (2022). DHARMa: residual diagnostics for hierarchical (multi-level/mixed) regression models. *R package version 0.4.6.1*.

Knauer, A. C., & Schiestl, F. P. (2017). The effect of pollinators and herbivores on selection for floral signals: a case study in Brassica rapa. *Evolutionary Ecology, 31*, 285-304. doi:10.1007/s10682-016-9878-8

Palacio, F. X., Ordano, M., & Benitez-Vieyra, S. (2019). Measuring natural selection on multivariate phenotypic traits: a protocol for verifiable and reproducible analyses of natural selection. *Israel Journal of Ecology and Evolution, 65*(3-4), 130-136. doi:10.1163/22244662-20191064

Rusman, Q., Traine, J., & Schiestl, F. P. (2025). Elevated temperature diminishes reciprocal selection in an experimental plant-pollinator-herbivore system. *Ecology Letters, 28*(1), e70060. doi:10.1111/ele.70060

Sahli, H. F., & Conner, J. K. (2011). Testing for conflicting and nonadditive selection: Floral adaptation to multiple pollinators through male and female fitness. *Evolution, 65*(5), 1457-1473. doi:10.1111/j.1558-5646.2011.01229.x

Stinchcombe, J. R., Agrawal, A. F., Hohenlohe, P. A., Arnold, S. J., & Blows, M. W. (2008). Estimating nonlinear selection gradients using quadratic regression coefficients: Double or nothing? *Evolution, 62*(9), 2435-2440. doi:10.1111/j.1558-5646.2008.00449.x

terHorst, C. P., Lau, J. A., Cooper, I. A., Keller, K. R., Rosa, R. J. L., Royer, A. M., . . . Conner, J. K. (2015). Quantifying nonadditive selection caused by indirect ecological effects. *Ecology, 96*(9), 2360-2369. doi:10.1890/14-0619.1

Therneau, T. M., & Lumley, T. (2015). Package 'survival'. *R Top Doc, 128*(10), 28-33.

Traine, J., Rusman, Q., & Schiestl, F. P. (2024). Too hot to handle: temperature-induced plasticity influences pollinator behaviour and plant fitness. *New Phytologist, 243*(4), 1571-1585. doi:10.1111/nph.19918

Wickham, H. (2011). ggplot2. *Wiley interdisciplinary reviews: computational statistics, 3*(2), 180-185.

**Table S1.** Results of experiments 1 and 2 showing plastic changes of *Brassica rapa* traits, insect behaviour, and plant reproduction in response to temperature. Output of (generalized) linear mixed models that tested effects of temperature (T), experiment (E), and their interaction (TxE) on the listed variables. Plants were grown in an ambient (23°C) or hot (27°C+ weekly 30°C day) temperature environment for 14 days prior to being measured, after which visitation by *Bombus terrestris* bumblebees and *Pieris rapae* butterflies occurred 2-4 days later (at 16-18 days). Bold values indicate results where $P \leq 0.05$.

| | Experiment 1 | | | | Experiment 2 | | | | | | | | | | | | |
|---|---|---|---|---|---|---|---|---|---|---|---|---|---|---|---|---|---|
| | Ambient | | Hot | | Ambient | | Hot | | T | | | E | | | TxE | | |
| | N | mean ± SE | N | mean ± SE | N | mean ± SE | N | mean ± SE | $\chi^2$ | df | P | $\chi^2$ | df | P | $\chi^2$ | df | P |
| Plant height (cm) | 292 | 37.30 ± 0.37 | 291 | 38.55 ± 0.37 | 97 | 30.96 ± 0.41 | 96 | 33.44 ± 0.64 | 13.98 | 1 | **<0.001** | 145.33 | 1 | **<0.001** | 2.67 | 1 | 0.103 |
| Leaf number | 292 | 9.75 ± 0.18 | 291 | 11.00 ± 0.24 | 97 | 5.12 ± 0.16 | 96 | 5.84 ± 0.16 | 24.19 | 1 | **< 0.001** | 347.48 | 1 | **<0.001** | 0.04 | 1 | 0.833 |
| Flower number | 292 | 12.00 ± 0.45 | 291 | 14.57 ± 0.43 | 97 | 11.94 ± 0.48 | 96 | 15.26 ± 0.81 | 33.01 | 1 | **< 0.001** | 0.21 | 1 | 0.646 | 2.27 | 1 | 0.131 |
| Nectar per flower (μl) | 287 | 0.08 ± 0.05 | 285 | 0.05 ± 0.04 | 97 | 0.11 ± 0.01 | 96 | 0.04 ± 0.00 | 87.75 | 1 | **<0.001** | 5.82 | 1 | **0.016** | 8.17 | 1 | **0.004** |
| Flower diameter (mm) | 287 | 13.22 ± 0.09 | 285 | 10.91 ± 0.08 | 97 | 15.04 ± 0.14 | 96 | 12.50 ± 0.12 | 581.81 | 1 | **< 0.001** | 228.39 | 1 | **<0.001** | 1.08 | 1 | 0.299 |
| Petal area ($cm^2$) | 286 | 0.25 ± 0.01 | 281 | 0.16 ± 0.01 | 97 | 0.22 ± 0.00 | 96 | 0.14 ± 0.00 | 1067.23 | 1 | **< 0.001** | 85.38 | 1 | **<0.001** | 0.24 | 1 | 0.622 |
| Petal length (cm) | 287 | 0.93 ± 0.01 | 281 | 0.79 ± 0.01 | 97 | 0.68 ± 0.01 | 96 | 0.57 ± 0.01 | 565.71 | 1 | **< 0.001** | 1639.86 | 1 | **<0.001** | 1.09 | 1 | 0.296 |
| Petal width (cm) | 287 | 0.46 ± 0.01 | 281 | 0.36 ± 0.01 | 97 | 0.55 ± 0.01 | 96 | 0.42 ± 0.01 | 918.31 | 1 | **< 0.001** | 287.17 | 1 | **<0.001** | 1.27 | 1 | 0.260 |
| Bumblebee plant visits | 195 | 1.77 ± 0.14 | 195 | 1.60 ± 0.14 | 97 | 1.07 ± 0.13 | 96 | 1.21 ± 0.13 | 0.17 | 1 | 0.684 | 13.98 | 1 | **<0.001** | 0.37 | 1 | 0.544 |
| Bumblebee flower visits | 195 | 19.77 ± 2.74 | 195 | 18.36 ± 3.12 | 97 | 15.00 ± 2.32 | 96 | 16.22 ± 2.40 | 0.02 | 1 | 0.881 | 8.96 | 1 | **0.003** | 0.04 | 1 | 0.846 |
| Butterfly plant visits | 192 | 63 ± 6 | 192 | 214 ± 13 | 49 | 50.92 ± 4.48 | 47 | 173.94 ± 10.43 | 167.19 | 1 | **<0.001** | 4.29 | 1 | **0.038** | 0.00 | 1 | 0.996 |
| Butterfly flower visits | 192 | 79 ± 8 | 192 | 255 ± 16 | 49 | 76.31 ± 6.81 | 47 | 223.36 ± 10.51 | 116.70 | 1 | **<0.001** | 0.52 | 1 | 0.470 | 0.22 | 1 | 0.642 |
| Number of fruits | 81 | 13 ± 1.9 | 81 | 11 ± 1.3 | 97 | 6.25 ± 0.76 | 96 | 6.19 ± 0.78 | 0.12 | 1 | 0.734 | 0.994 | 1 | 0.319 | 0.09 | 1 | 0.767 |
| Number of seeds | 81 | 129 ± 27 | 81 | 80 ± 14 | 97 | 44.68 ± 5.46 | 96 | 44.25 ± 5.60 | 1.46 | 1 | 0.228 | 0.09 | 1 | 0.759 | 1.27 | 1 | 0.259 |

**Table S2.** Results of experiment 1 and 2 showing the effect of temperature on linear selection differentials. Plants were visited by *Bombus terrestris* bumblebees in an ambient (23°C) or hot (27°C+ weekly 30°C day) temperature environment, where plants were previously kept for 16-18 days. Relative seed set was used as a fitness estimate in the models. Linear models regressing each trait against plant fitness were used to derive estimates. Because of zero-inflation in the fitness data of the *Bombus* treatment, non-parametric bootstraps were used to calculate standard errors of the gradients, and confidence intervals (C.I.) were used instead of p-values to assess significance. Bold values indicate results where $P \leq 0.05$ or the 95% C.I. does not include zero.

| | Experiment 1 | | | | Experiment 2 | | | |
|---|---|---|---|---|---|---|---|---|
| | Ambient | | Hot | | Ambient | | Hot | |
| | β ± SE | CI (95 %) | β ± SE | CI (95 %) | β ± SE | CI (95 %) | β ± SE | CI (95 %) |
| Plant height | **0.54 ± 0.24** | **0.168 – 1.148** | **0.45 ± 0.20** | **0.126 – 0.926** | **0.19 ± 0.11** | **0.143 – 0.429** | **0.46 ± 0.23** | **0.465 – 0.679** |
| Flower number | **0.71 ± 0.26** | **0.325 – 1.443** | 0.31 ± 0.26 | -0.076 – 1.036 | **0.22 ± 0.10** | **0.123 – 0.447** | 0.06 ± 0.06 | -0.112 – 0.181 |
| Leaf number | 0.46 ± 0.32 | -0.035 – 1.293 | 0.13 ± 0.24 | -0.284 – 0.646 | 0.15 ± 0.08 | -0.019 – 0.323 | 0.04 ± 0.07 | -0.117 – 0.222 |
| Flower diameter | -0.10 ± 0.21 | -0.675 – 0.228 | 0.16 ± 0.29 | -0.512 – 0.659 | -0.03 ± 0.07 | -0.188 – 0.136 | 0.08 ± 0.07 | -0.046 – 0.261 |
| Nectar per flower | **-0.29 ± 0.15** | **-0.662 – -0.038** | -0.05 ± 0.17 | -0.411 – 0.279 | **-0.24 ± 0.10** | **-0.457 – -0.162** | 0.04 ± 0.06 | -0.089 – 0.204 |
| Petal area | -0.24 ± 0.20 | -0.734 – 0.091 | 0.31 ± 0.19 | -0.034 – 0.708 | -0.01 ± 0.07 | -0.176 – 0.162 | 0.05 ± 0.07 | -0.091 – 0.244 |
| Petal length | -0.29 ± 0.26 | -0.939 – 0.127 | -0.04 ± 0.14 | -0.388 – 0.198 | -0.06 ± 0.07 | -0.224 – 0.107 | 0.03 ± 0.07 | -0.093 – 0.218 |
| Petal width | -0.09 ± 0.17 | -0.423 – 0.252 | 0.27 ± 0.19 | -0.074 – 0.677 | 0.03 ± 0.07 | -0.130 – 0.212 | 0.06 ± 0.08 | -0.112 – 0.249 |

**Table S3.** Plastic change of *Brassica rapa* traits in response to temperature. Mean trait values and output of (generalized) linear mixed models showing effects of elevated temperature on *Brassica rapa* plant traits. Plants were grown in an ambient (23°C) or hot (27°C+ weekly 30°C day) temperature environment for 14 days prior to being measured. Bold values indicate results where $P \leq 0.05$. Scent emission is expressed as pg / flower / l / h.

| | Ambient | | Hot | | Models | | |
|---|---|---|---|---|---|---|---|
| | N | mean ± SE | N | mean ± SE | $\chi^2$ | df | P |
| Days to flowering | 293 | 18.19 ± 0.06 | 293 | 16.70 ± 0.05 | 330.64 | 1 | **< 0.001** |
| Plant height (cm) | 292 | 37.30 ± 0.37 | 291 | 38.55 ± 0.37 | 9.38 | 1 | **0.002** |
| Leaf number | 292 | 9.75 ± 0.18 | 291 | 11.00 ± 0.24 | 21.99 | 1 | **< 0.001** |
| Flower number | 292 | 12.00 ± 0.45 | 291 | 14.57 ± 0.43 | 20.09 | 1 | **< 0.001** |
| Nectar per flower (µl) | 287 | 0.08 ± 0.05 | 285 | 0.05 ± 0.04 | 43.79 | 1 | **< 0.001** |
| Flower diameter (mm) | 287 | 13.22 ± 0.09 | 285 | 10.91 ± 0.08 | 472.94 | 1 | **< 0.001** |
| Petal area ($cm^2$) | 286 | 0.25 ± 0.01 | 281 | 0.16 ± 0.01 | 1069.18 | 1 | **< 0.001** |
| Petal length (cm) | 287 | 0.93 ± 0.01 | 281 | 0.79 ± 0.01 | 567.52 | 1 | **< 0.001** |
| Petal width (cm) | 287 | 0.46 ± 0.01 | 281 | 0.36 ± 0.01 | 893.59 | 1 | **< 0.001** |
| UV surface area | 285 | 0.28 ± 0.01 | 281 | 0.27 ± 0.01 | 2.62 | 1 | 0.105 |
| Benzaldehyde | 293 | 464.45 ± 28.65 | 282 | 567.53 ± 27.28 | 9.96 | 1 | **0.002** |
| 1-Butene-4-isothyocyanate | 293 | 188.50 ± 17.05 | 282 | 458.1 ± 29.55 | 138.21 | 1 | **< 0.001** |
| Methyl benzoate | 293 | 71.95 ± 5.38 | 282 | 35.26 ± 2.22 | 77.06 | 1 | **< 0.001** |
| Phenylethyl alcohol | 293 | 17.79 ± 2.27 | 282 | 7.10 ± 0.43 | 47.33 | 1 | **< 0.001** |
| 2-Amino benzaldehyde | 293 | 1312.34 ± 116.33 | 282 | 652.65 ± 59.03 | 22.71 | 1 | **< 0.001** |
| p-Anisaldehyde | 293 | 17.01 ± 1.76 | 282 | 7.38 ± 0.76 | 45.7 | 1 | **< 0.001** |
| Methyl anthranilate | 293 | 152.15 ± 12.64 | 282 | 53.81 ± 6.76 | 82 | 1 | **< 0.001** |
| *(Z)*-3-Hexen-1-ol acetate | 293 | 222.65 ± 17.58 | 282 | 150.67 ± 13.64 | 23.12 | 1 | **< 0.001** |
| Phenylacetaldehyde | 293 | 274.70 ± 49.89 | 282 | 53.86 ± 4.70 | 43.08 | 1 | **< 0.001** |
| Benzyl nitrile | 293 | 85.34 ± 6.58 | 282 | 18.60 ± 1.66 | 242.79 | 1 | **< 0.001** |
| Methyl salicylate | 293 | 87.11 ± 7.33 | 282 | 47.84 ± 3.14 | 18.4 | 1 | **< 0.001** |
| Indole | 293 | 240.80 ± 19.95 | 282 | 120.95 ± 10.23 | 26.75 | 1 | **< 0.001** |
| *(Z,Z)*-α-farnesene | 293 | 18.12 ± 1.10 | 282 | 10.70 ± 0.99 | 105.14 | 1 | **< 0.001** |
| *(E,E)*-α-farnesene | 293 | 367.04 ± 23.38 | 282 | 175.16 ± 9.74 | 138.49 | 1 | **< 0.001** |
| Total volatile emission | 293 | 3807.47 ± 223.42 | 282 | 2779.55 ± 125.78 | 13.99 | 1 | **< 0.001** |

**Table S4.** Changes in bumblebee and butterfly behaviour in response to temperature. Mean trait values and output of (generalized) linear mixed models showing effects of elevated temperature on the interactions of *Bombus terrestris* bumblebees and *Pieris rapae* butterflies with *Brassica rapa* plants. Visitation occurred in an ambient (23°C) or hot (27°C+ weekly 30°C day) temperature environment, where plants were previously kept for 16-18 days. Bold values indicate results where $P \leq 0.05$.

| | | | Ambient | Hot | model | | |
|---|---|---|---|---|---|---|---|
| | N | Trait | mean ± SE | mean ± SE | $\chi^2$ | df | P |
| *Bombus* | 195 | Flower visits | 19.77 ± 2.74 | 18.36 ± 3.12 | 0.13 | 1 | 0.723 |
| | 195 | Plant visits | 1.77 ± 0.14 | 1.60 ± 0.14 | 0.03 | 1 | 0.853 |
| | 195 | Seconds per plant | 129.67 ± 18.12 | 119.48 ± 21.17 | 3.45 | 1 | 0.063 |
| | 195 | Seconds per flower | 6.73 ± 0.46 | 6.47 ± 0.30 | 0.21 | 1 | 0.647 |
| *Bombus* in Generalized | 193 | Flower visits | 17.59 ± 2.39 | 20.54 ± 4.32 | 0.90 | 1 | 0.343 |
| | 193 | Plant visits | 1.11 ± 0.08 | 1.25 ± 0.09 | 0.00 | 1 | 0.963 |
| | 193 | Seconds per plant | 98.96 ± 15.07 | 114.63 ± 25.11 | 0.33 | 1 | 0.568 |
| | 193 | Seconds per flower | 5.83 ± 0.59 | 5.52 ± 0.30 | 0.33 | 1 | 0.564 |
| *Pieris** | 192 | Flower visits | 79 ± 8 | 255 ± 16 | 109.07 | 1 | **< 0.001** |
| | 192 | Plant visits | 63 ± 6 | 214 ± 13 | 84.89 | 1 | **< 0.001** |
| | 192 | Oviposition | 5.52 ± 0.40 | 8.95 ± 0.69 | 213.02 | 1 | **< 0.001** |
| *Pieris* in Generalized | 194 | Flower visits | 6.60 ± 0.76 | 34.27 ± 3.43 | 173.42 | 1 | **< 0.001** |
| | 194 | Plant visits | 4.97 ± 0.47 | 26.82 ± 2.41 | 194.02 | 1 | **< 0.001** |
| | 194 | Oviposition | 4.51 ± 0.30 | 7.66 ± 0.54 | 29.37 | 1 | **< 0.001** |

* Data from Rusman et al. (2025).

**Table S5.** Interactive effects of temperature and plant traits on pollinator visitation. Output of generalized linear mixed models testing the interactive effect of temperature on the relationship between pollinator flower visits and *Brassica rapa* plant traits. Plants were tested in an ambient (23°C) and hot (27°C+ weekly 30°C day) environment and visited by either *Pieris rapae* butterflies (*Pieris*), *Bombus terrestris* bumblebees (*Bombus*) or their combination (generalized). Bold values indicate results where $P \leq 0.05$. Italic values indicate results where $P \leq 0.1$.

| | *Bombus* | | | *Pieris* | | | Generalized | | |
|---|---|---|---|---|---|---|---|---|---|
| | $\chi^2$ | df | P | $\chi^2$ | df | P | $\chi^2$ | df | P |
| Plant height | 9.50 | 1 | **0.002** | 21.01 | 1 | **0.005** | 20.62 | 1 | **<0.001** |
| Temperature (T) | 1.09 | 1 | 0.297 | 112.65 | 1 | **<0.001** | 81.27 | 1 | **<0.001** |
| T x Plant height | 0.01 | 1 | 0.939 | 7.01 | 1 | **0.008** | 0.14 | 1 | 0.710 |
| Flower number | 32.71 | 1 | **<0.001** | 15.6 | 1 | *0.078* | 64.55 | 1 | **<0.001** |
| Temperature (T) | 0.12 | 1 | 0.728 | 63.74 | 1 | **<0.001** | 63.22 | 1 | **<0.001** |
| T x Flower number | 0.26 | 1 | 0.610 | 0.99 | 1 | 0.320 | 0.51 | 1 | 0.477 |
| Leaf number | 0.39 | 1 | 0.531 | 2.62 | 1 | 0.106 | 0.19 | 1 | 0.662 |
| Temperature (T) | 0.01 | 1 | 0.914 | 64.82 | 1 | **<0.001** | 80.29 | 1 | **<0.001** |
| T x Leaf number | 0.55 | 1 | 0.457 | 0.26 | 1 | 0.612 | 0.63 | 1 | 0.427 |
| Flower diameter | 6.22 | 1 | **0.013** | 10.53 | 1 | **0.001** | 16.91 | 1 | **<0.001** |
| Temperature (T) | 0.79 | 1 | 0.373 | 111.15 | 1 | **<0.001** | 99.6 | 1 | **<0.001** |
| T x Flower diameter | 0.47 | 1 | 0.492 | 0.43 | 1 | 0.514 | 0.41 | 1 | 0.524 |
| Nectar per flower | 0.26 | 1 | 0.613 | 0.07 | 1 | 0.795 | 0.07 | 1 | 0.789 |
| Temperature (T) | 0.06 | 1 | 0.800 | 95.1 | 1 | **<0.001** | 88.32 | 1 | **<0.001** |
| T x Nectar per flower | 6.75 | 1 | **0.009** | 0.21 | 1 | 0.647 | 6.7 | 1 | **0.010** |
| Petal area | 0.05 | 1 | 0.825 | 26.86 | 1 | **<0.001** | 8.13 | 1 | **0.004** |
| Temperature (T) | 0.02 | 1 | 0.876 | 120.13 | 1 | **<0.001** | 58.28 | 1 | **<0.001** |
| T x Petal area | 0.22 | 1 | 0.637 | 0.1 | 1 | 0.749 | 1.46 | 1 | 0.226 |
| Petal length | 0.2 | 1 | 0.651 | 8.33 | 1 | **0.004** | 4.15 | 1 | **0.042** |
| Temperature (T) | 0.33 | 1 | 0.564 | 101.7 | 1 | **<0.001** | 66.85 | 1 | **<0.001** |
| T x Petal length | 0.09 | 1 | 0.764 | 1.98 | 1 | 0.160 | 0 | 1 | 0.975 |
| Petal width | 0.05 | 1 | 0.824 | 28.11 | 1 | **<0.001** | 7.38 | 1 | **0.007** |
| Temperature (T) | 0.02 | 1 | 0.885 | 130.72 | 1 | **<0.001** | 64.81 | 1 | **<0.001** |
| T x Petal width | 0.05 | 1 | 0.829 | 0.24 | 1 | 0.624 | 2.91 | 1 | *0.088* |
| UV cover | 1.14 | 1 | 0.287 | 0.95 | 1 | 0.329 | 4.05 | 1 | **<0.001** |
| Temperature (T) | 0.28 | 1 | 0.598 | 118.62 | 1 | **<0.001** | 80.73 | 1 | **<0.001** |
| T x UV cover | 0.17 | 1 | 0.683 | 4.28 | 1 | **0.039** | 0.78 | 1 | 0.378 |
| Benzaldehyde | 1.2 | 1 | 0.273 | 0.08 | 1 | 0.782 | 15.55 | 1 | **<0.001** |
| Temperature (T) | 0.02 | 1 | 0.884 | 84.59 | 1 | **<0.001** | 114.13 | 1 | **<0.001** |
| T x Benzaldehyde | 0.38 | 1 | 0.536 | 0.7 | 1 | 0.404 | 1.21 | 1 | 0.270 |
| 1-Butene-4-isothyocyanate | 1.22 | 1 | 0.270 | 0 | 1 | 0.953 | 17.45 | 1 | **<0.001** |
| Temperature (T) | 0.02 | 1 | 0.886 | 65.52 | 1 | **<0.001** | 120.84 | 1 | **<0.001** |
| T x 1-butene-4-ITC | 0.01 | 1 | 0.923 | 0.02 | 1 | 0.894 | 0.76 | 1 | 0.384 |
| Methyl benzoate | 0.97 | 1 | 0.324 | 0.37 | 1 | 0.546 | 0.2 | 1 | 0.657 |
| Temperature (T) | 0 | 1 | 0.950 | 83.56 | 1 | **<0.001** | 75.46 | 1 | **<0.001** |
| T x methyl benzoate | 0.02 | 1 | 0.891 | 0.76 | 1 | 0.383 | 0.04 | 1 | 0.834 |
| Phenylethyl alcohol | 0.34 | 1 | 0.558 | 0.72 | 1 | 0.397 | 4.75 | 1 | **0.029** |
| Temperature (T) | 0.04 | 1 | 0.841 | 108.15 | 1 | **<0.001** | 78.1 | 1 | **<0.001** |
| T x phenylethyl alcohol | 0.33 | 1 | 0.566 | 0.34 | 1 | 0.558 | 0.01 | 1 | 0.918 |
| 2-Amino benzaldehyde | 2.85 | 1 | *0.091* | 0.18 | 1 | 0.670 | 0.32 | 1 | 0.573 |
| Temperature (T) | 0.34 | 1 | 0.562 | 79.21 | 1 | **<0.001** | 83.44 | 1 | **<0.001** |
| T x 2-amino benzaldehyde | 0.66 | 1 | 0.415 | 0.39 | 1 | 0.530 | 0.15 | 1 | 0.696 |
| p-Anisaldehyde | 0.02 | 1 | 0.895 | 0.21 | 1 | 0.650 | 0.47 | 1 | 0.493 |
| Temperature (T) | 0.08 | 1 | 0.775 | 80.7 | 1 | **<0.001** | 81.58 | 1 | **<0.001** |
| T x p-anisaldehyde | 2.26 | 1 | 0.133 | 2.11 | 1 | 0.146 | 0.46 | 1 | 0.499 |
| Methyl anthranilate | 2.04 | 1 | 0.153 | 0.24 | 1 | 0.625 | 0.34 | 1 | 0.560 |
| Temperature (T) | 0.02 | 1 | 0.884 | 74.11 | 1 | **<0.001** | 77.31 | 1 | **<0.001** |
| T x Methyl anthranilate | 0.1 | 1 | 0.750 | 0.78 | 1 | 0.377 | 0.04 | 1 | 0.841 |
| *(Z)*-3-Hexen-1-ol acetate | 6.5 | 1 | **0.011** | 1.82 | 1 | 0.178 | 10.97 | 1 | **0.001** |

| | | | | | | | | | |
|---|---|---|---|---|---|---|---|---|---|
| Temperature (T) | 1.03 | 1 | 0.310 | 88.96 | 1 | **<0.001** | 85.67 | 1 | **<0.001** |
| T x *(Z)*-3-hexen-1-ol acetate | 1.45 | 1 | 0.229 | 1.31 | 1 | 0.253 | 5.58 | 1 | **0.018** |
| Phenylacetaldehyde | 0.07 | 1 | 0.786 | 0.17 | 1 | 0.685 | 0.04 | 1 | 0.839 |
| Temperature (T) | 0.1 | 1 | 0.755 | 103.18 | 1 | **<0.001** | 73.7 | 1 | **<0.001** |
| T x phenylacetaldehyde | 0.43 | 1 | 0.512 | 0.22 | 1 | 0.638 | 1.65 | 1 | 0.199 |
| Benzyl nitrile | 2.25 | 1 | 0.133 | 1.06 | 1 | 0.303 | 0.25 | 1 | 0.617 |
| Temperature (T) | 0.09 | 1 | 0.766 | 75.01 | 1 | **<0.001** | 60.95 | 1 | **<0.001** |
| T x Benzyl nitrile | 2.49 | 1 | 0.115 | 1.53 | 1 | 0.216 | 0.84 | 1 | 0.361 |
| Methyl salicylate | 1.08 | 1 | 0.299 | 1.32 | 1 | 0.251 | 1.65 | 1 | 0.199 |
| Temperature (T) | 0.05 | 1 | 0.829 | 87.73 | 1 | **<0.001** | 81.89 | 1 | **<0.001** |
| T x methyl salicylate | 0.21 | 1 | 0.645 | 0.79 | 1 | 0.374 | 0 | 1 | 0.973 |
| Indole | 1.66 | 1 | 0.198 | 0.03 | 1 | 0.855 | 0.21 | 1 | 0.647 |
| Temperature (T) | 0.25 | 1 | 0.619 | 110.04 | 1 | **<0.001** | 76.8 | 1 | **<0.001** |
| T x indole | 0.45 | 1 | 0.501 | 0.2 | 1 | 0.655 | 0 | 1 | 0.969 |
| *(Z.Z)*-α-Farnesene | 2.21 | 1 | 0.137 | 2.83 | 1 | *0.092* | 1.8 | 1 | 0.180 |
| Temperature (T) | 0.09 | 1 | 0.759 | 124.81 | 1 | **<0.001** | 82.68 | 1 | **<0.001** |
| T x *(Z.Z)*-α-farnesene | 1.92 | 1 | 0.166 | 0.63 | 1 | 0.429 | 0.2 | 1 | 0.657 |
| *(E.E)*-α-Farnesene | 5.83 | 1 | **0.016** | 5.42 | 1 | **0.020** | 1.16 | 1 | 0.281 |
| Temperature (T) | 0.36 | 1 | 0.546 | 92.28 | 1 | **<0.001** | 84.97 | 1 | **<0.001** |
| T x *(E.E)*-α-farnesene | 0.6 | 1 | 0.438 | 0.44 | 1 | 0.507 | 0.13 | 1 | 0.714 |
| Total volatile emission | 0.01 | 1 | 0.934 | 0.23 | 1 | 0.632 | 3.55 | 1 | *0.060* |
| Temperature (T) | 0.13 | 1 | 0.722 | 83.17 | 1 | **<0.001** | 82.33 | 1 | **<0.001** |
| T x Total volatile emission | 2.91 | 1 | *0.088* | 0.76 | 1 | 0.385 | 0 | 1 | 0.988 |

**Table S6.** Interactive effects of temperature and plant traits on oviposition. Output of generalized linear mixed models testing the interactive effect of temperature on the relationship between butterfly oviposition and *Brassica rapa* plant traits. Plants were tested in an in an ambient (23°C) and hot (27°C+ weekly 30°C day) environment and visited by either *Pieris rapae* butterflies (*Pieris*) or by *Pieris rapae* butterflies and *Bombus terrestris* bumblebees (generalized). Bold values indicate results where $P \leq 0.05$. Italic values indicate results where $P \leq 0.1$.

| | *Pieris* | | | Generalized | | |
|---|---|---|---|---|---|---|
| | $\chi^2$ | df | P | $\chi^2$ | df | P |
| Plant height | 8.12 | 1 | **0.004** | 10.27 | 1 | **0.001** |
| Temperature (T) | 41.61 | 1 | **<0.001** | 28.04 | 1 | **<0.001** |
| T x plant height | 12.31 | 1 | **<0.001** | 2.94 | 1 | *0.086* |
| Flower number | 0.02 | 1 | 0.899 | 3.13 | 1 | *0.077* |
| Temperature (T) | 47.05 | 1 | **<0.001** | 26.11 | 1 | **<0.001** |
| T x flower number | 2.38 | 1 | 0.123 | 1.54 | 1 | 0.214 |
| Leaf number | 8.18 | 1 | **0.004** | 3.34 | 1 | *0.068* |
| Temperature (T) | 34.73 | 1 | **<0.001** | 19.13 | 1 | **<0.001** |
| T x leaf number | 3.93 | 1 | **0.047** | 0.64 | 1 | 0.424 |
| Flower diameter | 0.77 | 1 | 0.380 | 0.05 | 1 | 0.815 |
| Temperature (T) | 34.75 | 1 | **<0.001** | 14.92 | 1 | **<0.001** |
| T x flower diameter | 0 | 1 | 0.992 | 2.81 | 1 | *0.094* |
| Nectar per flower | 0.19 | 1 | 0.659 | 3.71 | 1 | *0.054* |
| Temperature (T) | 40.66 | 1 | **<0.001** | 32.34 | 1 | **<0.001** |
| T x nectar per flower | 0.05 | 1 | 0.824 | 0 | 1 | 0.980 |
| Petal area | 0.40 | 1 | 0.526 | 0.52 | 1 | 0.471 |
| Temperature (T) | 20.23 | 1 | **<0.001** | 7.63 | 1 | **0.006** |
| T x petal area | 0.01 | 1 | 0.935 | 3.28 | 1 | *0.070* |
| Petal length | 4.65 | 1 | **0.031** | 2.56 | 1 | 0.110 |
| Temperature (T) | 42.47 | 1 | **<0.001** | 24.52 | 1 | **<0.001** |
| T x petal length | 0.20 | 1 | 0.657 | 0.13 | 1 | 0.718 |
| UV cover | 0.05 | 1 | 0.828 | 1.49 | 1 | 0.223 |
| Temperature (T) | 19.02 | 1 | **<0.001** | 7.91 | 1 | **0.005** |
| T x UV cover | 0.13 | 1 | 0.714 | 2.74 | 1 | *0.098* |
| Benzaldehyde | 0.06 | 1 | 0.805 | 0.47 | 1 | 0.491 |

| | | | | | | |
|---|---|---|---|---|---|---|
| Temperature (T) | 50.7 | 1 | **<0.001** | 27.29 | 1 | **<0.001** |
| T x benzaldehyde | 2.12 | 1 | 0.146 | 0.42 | 1 | 0.517 |
| 1-Butene-4-isothyocyanate | 2.29 | 1 | 0.130 | 0.25 | 1 | 0.619 |
| Temperature (T) | 49.92 | 1 | **<0.001** | 28.41 | 1 | **<0.001** |
| T x 1-butene-4-isothyocyanate | 1.07 | 1 | 0.301 | 0 | 1 | 0.955 |
| Methyl benzoate | 0.49 | 1 | 0.486 | 1.94 | 1 | 0.164 |
| Temperature (T) | 46.21 | 1 | **<0.001** | 31.69 | 1 | **<0.001** |
| T x methyl benzoate | 2.40 | 1 | 0.121 | 7.77 | 1 | **<0.001** |
| Phenylethyl alcohol | 1.51 | 1 | 0.219 | 3.16 | 1 | *0.076* |
| Temperature (T) | 51.06 | 1 | **<0.001** | 32.81 | 1 | **<0.001** |
| T x phenylethyl alcohol | 1.58 | 1 | 0.209 | 0.41 | 1 | 0.524 |
| 2-Amino benzaldehyde | 0 | 1 | 0.987 | 0.20 | 1 | 0.656 |
| Temperature (T) | 44.55 | 1 | **<0.001** | 28.21 | 1 | **<0.001** |
| T x 2-Amino benzaldehyde | 0.05 | 1 | 0.832 | 2.58 | 1 | 0.108 |
| p-Anisaldehyde | 2.74 | 1 | 0.098 | 2.91 | 1 | *0.088* |
| Temperature (T) | 52.51 | 1 | **<0.001** | 32.52 | 1 | **<0.001** |
| T x p-anisaldehyde | 0.08 | 1 | 0.78 | 4.22 | 1 | **0.040** |
| Methyl anthranilate | 1.64 | 1 | 0.200 | 1.67 | 1 | 0.196 |
| Temperature (T) | 51.21 | 1 | **<0.001** | 31.92 | 1 | **<0.001** |
| T x methyl anthranilate | 0.59 | 1 | 0.441 | 6.21 | 1 | **0.013** |
| *(Z)*-3-Hexen-1-ol acetate | 1.43 | 1 | 0.231 | 0.95 | 1 | 0.331 |
| Temperature (T) | 40.35 | 1 | **<0.001** | 28.04 | 1 | **<0.001** |
| T x *(Z)*-3-hexen-1-ol acetate | 2.28 | 1 | 0.131 | 8.51 | 1 | **0.004** |
| Phenylacetaldehyde | 1.23 | 1 | 0.268 | 0.02 | 1 | 0.898 |
| Temperature (T) | 51.00 | 1 | **<0.001** | 28.00 | 1 | **<0.001** |
| T x phenylacetaldehyde | 2.44 | 1 | 0.118 | 0 | 1 | 0.978 |
| Benzyl nitrile | 0.18 | 1 | 0.672 | 0.14 | 1 | 0.708 |
| Temperature (T) | 43.78 | 1 | **<0.001** | 20.86 | 1 | **<0.001** |
| T x benzyl nitrile | 0.09 | 1 | 0.767 | 3.02 | 1 | *0.082* |
| Methyl salicylate | 2.17 | 1 | 0.141 | 0.99 | 1 | 0.320 |
| Temperature (T) | 47.34 | 1 | **<0.001** | 24.22 | 1 | **<0.001** |
| T x methyl salicylate | 0.27 | 1 | 0.600 | 4.41 | 1 | 0.036 |
| Indole | 0.28 | 1 | 0.596 | 6.57 | 1 | **0.010** |
| Temperature (T) | 49.76 | 1 | **<0.001** | 35.94 | 1 | **<0.001** |
| T x indole | 0.66 | 1 | 0.416 | 0.67 | 1 | 0.415 |
| *(Z.Z)*-α-Farnesene | 1.13 | 1 | 0.289 | 4.30 | 1 | **0.038** |
| Temperature (T) | 44.99 | 1 | **<0.001** | 34.71 | 1 | **<0.001** |
| T x *(Z.Z)*-α-farnesene | 0.87 | 1 | 0.350 | 5.85 | 1 | **0.016** |
| *(E.E)*-α-Farnesene | 2.62 | 1 | 0.106 | 7.73 | 1 | **0.005** |
| Temperature (T) | 54.52 | 1 | **<0.001** | 37.57 | 1 | **<0.001** |
| T x *(E.E)*-α-farnesene | 5.27 | 1 | **0.022** | 0.03 | 1 | 0.864 |
| Total volatile emission | 4.59 | 1 | **0.032** | 5.42 | 1 | **0.020** |
| Temperature (T) | 54.38 | 1 | **<0.001** | 35.35 | 1 | **<0.001** |
| T x Total volatile emission | 1.07 | 1 | 0.301 | 0.43 | 1 | 0.513 |

**Table S7.** Effects of plant traits on pollinator visitation in different temperature environments. Output of generalized linear mixed models testing the relationship between pollinator flower visits and *Brassica rapa* plant traits in an ambient (23°C) and hot (27°C+ weekly 30°C day) environment and when plants were visited by *Bombus terrestris* bumblebees (*Bombus*), *Pieris rapae* butterflies (*Pieris*), or their combination (generalized treatment). Bold values indicate results where *P* ≤ 0.05. Italic values indicate results where *P* ≤ 0.1.

| | *Bombus* | | | | | | *Pieris* | | | | | | Generalized | | | | | |
|---|---|---|---|---|---|---|---|---|---|---|---|---|---|---|---|---|---|---|
| | Ambient | | | Hot | | | Ambient | | | Hot | | | Ambient | | | Hot | | |
| | $\chi^2$ | df | P | $\chi^2$ | df | P | $\chi^2$ | df | P | $\chi^2$ | df | P | $\chi^2$ | df | P | $\chi^2$ | df | P |
| Plant height | 9.34 | 1 | **0.002** | 7.82 | 1 | **0.005** | 30.48 | 1 | **< 0.001** | 3.34 | 1 | 0.068 | 4.39 | 1 | **0.036** | 26.65 | 1 | **< 0.001** |
| Flower number | 21.73 | 1 | **< 0.001** | 19.26 | 1 | **< 0.001** | 10.78 | 1 | **0.001** | 10.42 | 1 | **0.001** | 23.85 | 1 | **< 0.001** | 43.96 | 1 | **< 0.001** |
| Leaf number | 6.86 | 1 | **0.009** | 0.03 | 1 | 0.867 | 2.10 | 1 | 0.147 | 1.98 | 1 | 0.160 | 1.04 | 1 | 0.308 | 0.78 | 1 | 0.376 |
| Flower diameter | 0.11 | 1 | 0.743 | 8.83 | 1 | **0.003** | 6.25 | 1 | **0.012** | 3.13 | 1 | *0.077* | 4.53 | 1 | **0.033** | 15.54 | 1 | **< 0.001** |
| Nectar per flower | 0.06 | 1 | 0.808 | 1.10 | 1 | 0.295 | 1.00 | 1 | 0.317 | 0.00 | 1 | 0.994 | *3.58* | 1 | *0.058* | 7.83 | 1 | **0.005** |
| Petal area | 0.37 | 1 | 0.545 | 1.39 | 1 | 0.238 | 12.53 | 1 | **< 0.001** | 9.86 | 1 | **0.002** | 2.66 | 1 | 0.103 | 14.05 | 1 | **< 0.001** |
| Petal length | 0.36 | 1 | 0.546 | 0.12 | 1 | 0.725 | 6.76 | 1 | **0.009** | 1.31 | 1 | 0.253 | 2.93 | 1 | *0.087* | 2.44 | 1 | 0.118 |
| Petal width | 0.11 | 1 | 0.738 | 0 | 1 | 0.984 | 14.15 | 1 | **<0.001** | 14.17 | 1 | **<0.001** | 1.41 | 1 | 0.235 | 18.21 | 1 | **<0.001** |
| UV cover | 1.56 | 1 | 0.211 | 0.32 | 1 | 0.574 | 4.99 | 1 | **0.026** | 0.81 | 1 | 0.369 | 1.42 | 1 | 0.233 | 6.88 | 1 | **0.009** |
| Benzaldehyde | 0.04 | 1 | 0.851 | 2.56 | 1 | 0.110 | 0.46 | 1 | 0.499 | 0.24 | 1 | 0.621 | 12.73 | 1 | **< 0.001** | 4.53 | 1 | **0.033** |
| 1-Butene-4-isothyocyanate | 0.01 | 1 | 0.905 | 0.14 | 1 | 0.710 | 0.01 | 1 | 0.907 | 0.01 | 1 | 0.933 | 10.28 | 1 | **0.001** | 6.33 | 1 | **0.012** |
| Methyl benzoate | 0.44 | 1 | 0.505 | 0.02 | 1 | 0.898 | 0.91 | 1 | 0.339 | 0.11 | 1 | 0.744 | 0.27 | 1 | 0.604 | 0.45 | 1 | 0.501 |
| Phenylethyl alcohol | 1.92 | 1 | 0.166 | 0.19 | 1 | 0.661 | 0.37 | 1 | 0.542 | 0.32 | 1 | 0.572 | 2.65 | 1 | 0.103 | 2.88 | 1 | *0.089* |
| 2-Amino benzaldehyde | 3.83 | 1 | **0.050** | 0.56 | 1 | 0.455 | 0.48 | 1 | 0.487 | 0.04 | 1 | 0.833 | 0.08 | 1 | 0.772 | 0.01 | 1 | 0.912 |
| p-Anisaldehyde | 2.07 | 1 | 0.150 | *3.70* | 1 | *0.054* | 1.36 | 1 | 0.243 | 1.89 | 1 | 0.169 | 0.19 | 1 | 0.659 | 0.32 | 1 | 0.573 |
| Methyl anthranilate | 2.58 | 1 | 0.108 | 0.13 | 1 | 0.717 | 0.88 | 1 | 0.349 | 0.24 | 1 | 0.621 | 0.00 | 1 | 0.965 | 0.48 | 1 | 0.490 |
| *(Z)*-3-Hexen-1-ol acetate | 1.73 | 1 | 0.189 | 4.82 | 1 | **0.028** | 2.27 | 1 | 0.132 | 0.16 | 1 | 0.687 | 13.87 | 1 | **< 0.001** | 0.46 | 1 | 0.497 |
| Phenylacetaldehyde | 0.59 | 1 | 0.442 | 0.23 | 1 | 0.628 | 0.45 | 1 | 0.505 | 0.32 | 1 | 0.571 | 0.10 | 1 | 0.749 | *3.64* | 1 | *0.056* |
| Benzyl nitrile | 5.30 | 1 | **0.021** | 0.14 | 1 | 0.709 | 2.18 | 1 | 0.140 | 0.17 | 1 | 0.678 | 0.08 | 1 | 0.778 | 0.01 | 1 | 0.930 |
| Methyl salicylate | 2.53 | 1 | 0.112 | 0.04 | 1 | 0.845 | 1.90 | 1 | 0.168 | 0.00 | 1 | 0.977 | 1.36 | 1 | 0.243 | 1.47 | 1 | 0.225 |
| Indole | 3.06 | 1 | *0.080* | 0.21 | 1 | 0.645 | 0.70 | 1 | 0.403 | 0.32 | 1 | 0.574 | 0.65 | 1 | 0.419 | 0.36 | 1 | 0.547 |
| *(Z.Z)*-α-farnesene | 6.54 | 1 | **0.011** | 0.72 | 1 | 0.398 | 0.46 | 1 | 0.498 | 2.25 | 1 | 0.133 | 0.01 | 1 | 0.903 | 0.47 | 1 | 0.494 |
| *(E.E)*-α-farnesene | 7.92 | 1 | **0.005** | 2.11 | 1 | 0.147 | 4.22 | 1 | **0.040** | 2.74 | 1 | *0.098* | 0.28 | 1 | 0.597 | 0.26 | 1 | 0.609 |
| Total volatile emission | 3.26 | 1 | *0.071* | 0.29 | 1 | 0.591 | 0.75 | 1 | 0.385 | 0.23 | 1 | 0.635 | 2.55 | 1 | 0.110 | 1.21 | 1 | 0.272 |

**Table S8.** Effects of plant traits on oviposition in different temperature environments. Output of generalized linear mixed models testing the relationship between butterfly oviposition per plant and *Brassica rapa* plant traits in an ambient (23°C) and hot (27°C+ weekly 30°C day) environment and when plants were visited by either *Pieris rapae* butterflies (*Pieris*) or by *Pieris rapae* butterflies and *Bombus terrestris* bumblebees (generalized). Bold values indicate results where $P \leq 0.05$. Italic values indicate results where $P \leq 0.1$.

| | *Pieris* | | | | | | Generalized | | | | | |
|---|---|---|---|---|---|---|---|---|---|---|---|---|
| | Ambient | | | Hot | | | Ambient | | | Hot | | |
| | $\chi^2$ | df | P | $\chi^2$ | df | P | $\chi^2$ | df | P | $\chi^2$ | df | P |
| Plant height | 18.90 | 1 | **< 0.001** | 0.05 | 1 | 0.823 | 7.14 | 1 | **0.008** | 2.22 | 1 | 0.136 |
| Flower number | 0.04 | 1 | 0.838 | 1.99 | 1 | 0.158 | 3.73 | 1 | *0.053* | 0.02 | 1 | 0.890 |
| Leaf number | 4.06 | 1 | **0.044** | 2.37 | 1 | 0.124 | 0.71 | 1 | 0.398 | 0.66 | 1 | 0.416 |
| Flower diameter | 0.87 | 1 | 0.350 | 0.34 | 1 | 0.562 | 1.67 | 1 | 0.196 | 0.85 | 1 | 0.356 |
| Nectar per flower | 0.54 | 1 | 0.462 | < 0.01 | 1 | 0.951 | 0.32 | 1 | 0.573 | 1.61 | 1 | 0.205 |
| Petal area | 1.18 | 1 | 0.277 | 0.13 | 1 | 0.716 | 5.38 | 1 | **0.020** | 0.61 | 1 | 0.435 |
| Petal length | 2.71 | 1 | 0.100 | *3.17* | 1 | *0.075* | 0.17 | 1 | 0.681 | 2.25 | 1 | 0.133 |
| Petal width | 1.63 | 1 | 0.202 | 0.06 | 1 | 0.807 | 7.62 | 1 | **0.006** | 0.10 | 1 | 0.751 |
| UV cover | 0.17 | 1 | 0.677 | 0.57 | 1 | 0.452 | 0.01 | 1 | 0.926 | 0.76 | 1 | 0.384 |
| Benzaldehyde | 2.69 | 1 | 0.101 | < 0.01 | 1 | 0.994 | 3.44 | 1 | *0.064* | 0.16 | 1 | 0.692 |
| 1-Butene-4-isothyocyanate | 0.22 | 1 | 0.636 | < 0.01 | 1 | 0.965 | 0.10 | 1 | 0.751 | 7.27 | 1 | **0.007** |
| Methyl benzoate | 1.96 | 1 | 0.162 | 0.20 | 1 | 0.656 | 0.21 | 1 | 0.645 | 0.51 | 1 | 0.475 |
| Phenylethyl alcohol | 0.09 | 1 | 0.764 | 0.02 | 1 | 0.893 | 0.24 | 1 | 0.623 | 0.87 | 1 | 0.351 |
| 2-Amino benzaldehyde | 0.24 | 1 | 0.623 | 0.51 | 1 | 0.473 | 3.49 | 1 | *0.062* | 0.01 | 1 | 0.943 |
| p-Anisaldehyde | 0.78 | 1 | 0.377 | 0.01 | 1 | 0.936 | 8.99 | 1 | **0.003** | 0.83 | 1 | 0.363 |
| Methyl anthranilate | 1.13 | 1 | 0.287 | 0.04 | 1 | 0.838 | 9.42 | 1 | **0.002** | 1.80 | 1 | 0.180 |
| *(Z)*-3-Hexen-1-ol acetate | 4.91 | 1 | **0.027** | 0.10 | 1 | 0.755 | 1.25 | 1 | 0.264 | 0.01 | 1 | 0.905 |
| Phenylacetaldehyde | 0.13 | 1 | 0.723 | < 0.01 | 1 | 0.971 | 1.16 | 1 | 0.282 | 2.56 | 1 | 0.109 |
| Benzyl nitrile | 1.21 | 1 | 0.270 | 0.14 | 1 | 0.711 | 0.67 | 1 | 0.413 | 0.66 | 1 | 0.417 |
| Methyl salicylate | 0.28 | 1 | 0.594 | 0.14 | 1 | 0.704 | 2.14 | 1 | 0.144 | 1.35 | 1 | 0.246 |
| Indole | 0.21 | 1 | 0.644 | 0.18 | 1 | 0.669 | 13.34 | 1 | **< 0.001** | 0.05 | 1 | 0.819 |
| *(Z.Z)*-α-farnesene | 0.40 | 1 | 0.528 | 0.27 | 1 | 0.606 | 0.29 | 1 | 0.591 | 5.34 | 1 | **0.021** |
| *(E.E)*-α-farnesene | 2.15 | 1 | 0.143 | 1.44 | 1 | 0.230 | 0.86 | 1 | 0.353 | 1.71 | 1 | 0.191 |
| Total volatile emission | 0.93 | 1 | 0.336 | < 0.01 | 1 | 0.960 | 7.90 | 1 | **0.005** | 0.88 | 1 | 0.348 |

**Table S9.** The interactive effects of temperature and biotic treatments on the directional selection of plant traits. Directional selection gradients on plant visual traits and flower scent compounds under the interactive effects of temperature environment and visits by either *Bombus terrestris* bumblebees (*Bombus*), *Pieris rapae* butterflies (*Pieris*), or their combination (generalized). Visitation occurred in in an ambient (23°C) or hot (27°C+ weekly 30°C day) temperature environment. where plants were previously kept for 16-18 days. Relative seed set was used as a fitness estimate in the models. Selection for morphological and scent traits was analysed separately. Because of zero-inflation in the fitness data of the *Bombus* treatment. non-parametric bootstraps were used to calculate standard errors of the gradients. and confidence intervals (C.I.) were used instead of p-values to assess significance. Bold values indicate results where $P \leq 0.05$ or the 95% C.I. does not include zero. Italic values indicate results where $P \leq 0.1$.

| | *Bombus* | | | *Pieris* | | | Generalized | | |
|---|---|---|---|---|---|---|---|---|---|
| | β ± SE | CI 95% | VIF | β ± SE | P | VIF | β ± SE | P | VIF |
| Temperature (T) | 0.17 ± 0.46 | -0.766 – 1.057 | - | -0.05 ± 0.17 | 0.966 | - | -0.14 ± 0.17 | 0.291 | - |
| Flowering time | 0.18 ± 0.22 | -0.240 – 0.600 | 1.50 | -0.05 ± 0.08 | 0.189 | 1.54 | 0.13 ± 0.10 | 0.963 | 1.91 |
| Plant height | 0.44 ± 0.26 | -0.054 – 0.994 | 1.24 | **0.19 ± 0.07** | **0.001** | 1.25 | **0.16 ± 0.07** | **0.021** | 1.18 |
| Flower number | **0.62 ± 0.23** | **0.250 – 1.188** | 1.19 | **0.30 ± 0.07** | **< 0.001** | 1.32 | **0.23 ± 0.06** | **< 0.001** | 1.25 |
| Leaf number | -0.11 ± 0.30 | -0.604 – 0.589 | 1.14 | -0.11 ± 0.08 | 0.330 | 1.30 | *-0.03 ± 0.07* | *0.072* | 1.32 |
| Flower diameter | -0.13 ± 0.29 | -0.813 – 0.368 | 2.76 | -0.02 ± 0.11 | 0.672 | 3.29 | *0.16 ± 0.11* | *0.067* | 3.16 |
| Nectar per flower | **-0.31 ± 0.14** | **-0.623 – -0.060** | 1.07 | -0.06 ± 0.05 | 0.565 | 1.28 | -0.07 ± 0.06 | 0.669 | 1.12 |
| Petal area | -0.03 ± 0.28 | -0.539 – 0.598 | 3.30 | 0.15 ± 0.13 | 0.145 | 4.25 | -0.24 ± 0.12 | 0.147 | 4.02 |
| UV cover | 0.09 ± 0.29 | -0.346 – 0.830 | 1.05 | 0.06 ± 0.06 | 0.998 | 1.08 | 0.01 ± 0.06 | 0.745 | 1.13 |
| T X Flowering time | -0.12 ± 0.40 | -0.953 – 0.610 | - | -0.05 ± 0.11 | 0.667 | - | *-0.23 ± 0.13* | *0.082* | - |
| T X Plant height | -0.11 ± 0.33 | -0.759 – 0.581 | - | -0.05 ± 0.09 | 0.614 | - | -0.09 ± 0.10 | 0.337 | - |
| T X Flower number | -0.19 ± 0.37 | -0.853 – 0.629 | - | -0.03 ± 0.09 | 0.757 | - | -0.11 ± 0.10 | 0.264 | - |
| T X Leaf number | 0.32 ± 0.42 | -0.558 – 1.096 | - | **0.23 ± 0.10** | **0.026** | - | **0.23 ± 0.10** | **0.021** | - |
| T X Flower diameter | -0.06 ± 0.69 | -1.740 – 1.028 | - | -0.01 ± 0.15 | 0.941 | - | -0.04 ± 0.16 | 0.776 | - |
| T X Nectar per flower | 0.11 ± 0.33 | -0.509 – 0.820 | - | 0.15 ± 0.11 | 0.189 | - | 0.12 ± 0.09 | 0.219 | - |
| T X Petal area | 0.70 ± 0.51 | -0.185 – 1.856 | - | -0.01 ± 0.20 | 0.968 | - | 0.24 ± 0.20 | 0.228 | - |
| T X UV cover | 0.07 ± 0.32 | -0.699 – 0.606 | - | -0.12 ± 0.09 | 0.148 | - | -0.06 ± 0.09 | 0.494 | - |
| Temperature (T) | 0.38 ± 0.52 | -0.592 – 1.465 | - | 0.16 ± 0.17 | 0.421 | - | **0.47 ± 0.15** | **0.008** | - |
| Benzaldehyde | -0.38 ± 0.37 | -1.194 – 0.269 | 2.08 | 0.00 ± 0.16 | 0.657 | 5.12 | **-0.41 ± 0.10** | **0.001** | 2.59 |
| 1-Butene-4-isothyocyanate | 0.25 ± 0.25 | -0.223 – 0.773 | 1.45 | -0.10 ± 0.12 | 0.874 | 2.43 | -0.04 ± 0.10 | 0.111 | 1.80 |
| Methyl benzoate | 0.01 ± 0.24 | -0.455 – 0.483 | 1.48 | -0.03 ± 0.16 | 0.859 | 5.71 | *0.17 ± 0.11* | *0.073* | 2.99 |
| Phenylethyl alcohol | 0.68 ± 0.57 | -0.356 – 1.923 | 7.27 | -0.02 ± 0.17 | 0.354 | 5.36 | 0.04 ± 0.14 | 0.731 | 4.83 |
| 2-Amino benzaldehyde | -0.42 ± 0.51 | -1.689 – 0.399 | 6.44 | 0.10 ± 0.20 | 0.207 | 6.86 | 0.18 ± 0.15 | 0.672 | 5.07 |
| p-Anisaldehyde | 0.41 ± 0.35 | -0.166 – 1.218 | 1.87 | 0.08 ± 0.12 | 0.351 | 2.95 | 0.02 ± 0.11 | 0.474 | 2.39 |
| Methyl anthranilate | 0.24 ± 0.38 | -0.554 – 0.924 | 3.58 | 0.06 ± 0.17 | 0.613 | 6.06 | -0.10 ± 0.15 | 0.800 | 6.09 |
| *(Z)*-3-Hexen-1-ol acetate | -0.01 ± 0.21 | -0.450 – 0.395 | 1.39 | 0.03 ± 0.10 | 0.965 | 2.00 | 0.15 ± 0.09 | 0.101 | 1.74 |
| Phenylacetaldehyde | **-1.27 ± 0.65** | **-2.778 – -0.176** | 6.96 | -0.25 ± 0.18 | 0.392 | 6.06 | -0.15 ± 0.12 | 0.435 | 3.92 |
| Benzyl nitrile | 0.53 ± 0.70 | -0.636 – 2.079 | 4.17 | 0.04 ± 0.23 | 0.744 | 7.53 | 0.17 ± 0.16 | 0.232 | 5.29 |
| Methyl salicylate | 0.07 ± 0.23 | -0.412 – 0.497 | 1.47 | 0.00 ± 0.16 | 0.691 | 4.27 | -0.03 ± 0.10 | 0.410 | 2.84 |
| Indole | -0.28 ± 0.50 | -1.342 – 0.644 | 5.97 | -0.04 ± 0.27 | 0.643 | 9.31 | -0.11 ± 0.17 | 0.672 | 6.82 |
| *(Z.Z)*-α-Farnesene | 0.39 ± 0.66 | -0.765 – 1.877 | 9.04 | -0.16 ± 0.11 | 0.076 | 1.83 | 0.19 ± 0.16 | 0.297 | 7.56 |
| *(E.E)*-α-Farnesene | 0.04 ± 0.58 | -1.268 – 1.045 | 8.09 | **0.32 ± 0.14** | **0.004** | 3.75 | 0.05 ± 0.15 | 0.694 | 6.94 |
| T X benzaldehyde | 1.00 ± 0.60 | -0.125 – 2.235 | - | -0.11 ± 0.24 | 0.646 | - | **0.35 ± 0.16** | **0.032** | - |
| T X 1-butene-4-isothyocyanate | -0.26 ± 0.37 | -1.004 – 0.441 | - | 0.16 ± 0.17 | 0.330 | - | -0.12 ± 0.14 | 0.369 | - |
| T X methyl benzoate | 0.16 ± 0.57 | -0.868 – 1.377 | - | 0.12 ± 0.25 | 0.625 | - | -0.05 ± 0.17 | 0.781 | - |
| T X phenylethyl alcohol | -0.34 ± 0.84 | -2.037 – 1.254 | - | -0.18 ± 0.25 | 0.457 | - | -0.16 ± 0.21 | 0.440 | - |
| T X 2-amino benzaldehyde | 1.29 ± 0.93 | -0.343 – 3.402 | - | 0.12 ± 0.26 | 0.658 | - | -0.28 ± 0.22 | 0.203 | - |
| T X p-anisaldehyde | -0.54 ± 0.46 | -1.478 – 0.308 | - | 0.00 ± 0.18 | 0.985 | - | 0.08 ± 0.15 | 0.625 | - |
| T X methyl anthranilate | -0.29 ± 0.60 | -1.445 – 0.995 | - | -0.24 ± 0.25 | 0.334 | - | 0.19 ± 0.23 | 0.415 | - |
| T X *(Z)*-3-hexen-1-ol acetate | -0.24 ± 0.45 | -1.158 – 0.640 | - | -0.06 ± 0.14 | 0.657 | - | -0.07 ± 0.13 | 0.562 | - |
| T X phenylacetaldehyde | -0.21 ± 1.06 | -2.362 – 1.779 | - | 0.32 ± 0.30 | 0.275 | - | 0.25 ± 0.21 | 0.245 | - |
| T X benzyl nitrile | 0.16 ± 0.81 | -1.623 – 1.587 | - | 0.02 ± 0.30 | 0.949 | - | -0.06 ± 0.24 | 0.813 | - |
| T X methyl salicylate | -0.13 ± 0.39 | -0.870 – 0.637 | - | -0.09 ± 0.22 | 0.691 | - | -0.09 ± 0.16 | 0.572 | - |
| T X indole | -0.58 ± 1.00 | -2.833 – 1.209 | - | -0.06 ± 0.34 | 0.871 | - | 0.13 ± 0.25 | 0.603 | - |
| T X *(Z.Z)*-α-farnesene | -1.08 ± 0.96 | -3.240 – 0.568 | - | 0.05 ± 0.14 | 0.736 | - | -0.16 ± 0.28 | 0.570 | - |
| T X *(E.E)*-α-farnesene | 0.70 ± 0.92 | -0.939 – 2.706 | - | -0.07 ± 0.19 | 0.721 | - | -0.02 ± 0.26 | 0.936 | - |

**Table S10.** The effects of temperature on the directional selection of plant traits under different biotic treatments. Directional selection gradients on plant visual traits and flower scent compounds for plants grown and tested in two temperature environments and visited by either *Bombus terrestris* bumblebees (*Bombus*), *Pieris rapae* butterflies (*Pieris*), or their combination (generalized). Visitation occurred in in an ambient (23ºC) or hot (27ºC+ weekly 30ºC day) temperature environment. where plants were previously kept for 16-18 days. Relative seed set was used as a fitness estimate in the models. Selection for morphological and scent traits were analysed separately. Because of zero-inflation in the fitness data of the *Bombus* treatment. non-parametric bootstraps were used to calculate standard errors of the gradients. and confidence intervals (C.I.) were used instead of p-values to assess significance. Bold values indicate results where *P* ≤ 0.05 or the 95% C.I. does not include zero. Italic values indicate results where *P* ≤ 0.1 or the 90% C.I. does not include zero. 1-Butene-4-ITC = 1-butene-4-isothiocyanate. 3-hexenyl acetate = *(Z)*-3-Hexen-1-ol acetate.

| | *Bombus* | | | | | | | | *Pieris* | | | | | | Generalized | | | | | |
|---|---|---|---|---|---|---|---|---|---|---|---|---|---|---|---|---|---|---|---|---|
| | Ambient | | | | Hot | | | | Ambient | | | Hot | | | Ambient | | | Hot | | |
| | β ± SE | CI (95 %) | CI (90 %) | VIF | β ± SE | CI (95 %) | CI (90 %) | VIF | β ± SE | P | VIF | β ± SE | P | VIF | β ± SE | P | VIF | β ± SE | P | VIF |
| Flowering time | 0.15 ± 0.18 | -0.189 – 0.515 | -0.136 – 0.453 | 1.11 | 0.04 ± 0.22 | -0.449 – 0.421 | -0.364 – 0.362 | 1.06 | -0.04 ± 0.07 | 0.580 | 1.23 | -0.07 ± 0.05 | 0.143 | 1.13 | 0.10 ± 0.08 | 0.254 | 1.25 | -0.07 ± 0.05 | 0.178 | 1.12 |
| Plant height | *0.42 ± 0.25* | *-0.039 – 0.956* | *0.031 – 0.872* | 1.39 | *0.34 ± 0.21* | *-0.016 – 0.814* | *0.043 – 0.734* | 1.21 | **0.19 ± 0.08** | **0.018** | 1.55 | **0.14 ± 0.05** | **0.007** | 1.13 | *0.15 ± 0.08* | *0.052* | 1.25 | 0.08 ± 0.06 | 0.187 | 1.30 |
| Flower number | **0.66 ± 0.25** | **0.279 – 1.281** | 0.335 – 1.167 | 1.57 | *0.40 ± 0.27* | *-0.005 – 1.113* | *0.047 – 0.983* | 1.27 | **0.28 ± 0.08** | **<.001** | 1.47 | **0.26 ± 0.05** | **<.001** | 1.11 | **0.25 ± 0.07** | **0.001** | 1.19 | *0.10 ± 0.06* | *0.083* | 1.38 |
| Leaf number | -0.11 ± 0.28 | -0.574 – 0.582 | -0.508 – 0.450 | 1.67 | 0.20 ± 0.30 | -0.407 – 0.764 | -0.315 – 0.668 | 1.40 | -0.09 ± 0.08 | 0.274 | 1.6 | **0.13 ± 0.05** | **0.014** | 1.15 | -0.03 ± 0.07 | 0.691 | 1.14 | **0.19 ± 0.06** | **0.001** | 1.24 |
| Flower diameter | -0.13 ± 0.28 | -0.769 – 0.353 | -0.641 – 0.282 | 2.17 | -0.12 ± 0.40 | -1.165 – 0.481 | -0.984 – 0.424 | 1.84 | -0.02 ± 0.09 | 0.847 | 2.2 | -0.03 ± 0.06 | 0.658 | 1.57 | 0.12 ± 0.09 | 0.176 | 1.61 | 0.09 ± 0.07 | 0.233 | 2.12 |
| Nectar per flower | **-0.34 ± 0.16** | **-0.700 – -0.071** | -0.635 – -0.115 | 1.09 | -0.18 ± 0.26 | -0.669 – 0.362 | -0.590 – 0.268 | 1.18 | -0.07 ± 0.07 | 0.320 | 1.13 | 0.05 ± 0.05 | 0.295 | 1.19 | -0.07 ± 0.07 | 0.321 | 1.12 | 0.04 ± 0.05 | 0.450 | 1.10 |
| Petal area | -0.03 ± 0.23 | -0.438 – 0.493 | -0.369 – 0.397 | 2.30 | **0.44 ± 0.27** | **0.034 – 1.142** | 0.096 – 1.006 | 1.84 | 0.10 ± 0.10 | 0.316 | 2.23 | 0.07 ± 0.06 | 0.280 | 1.71 | -0.16 ± 0.09 | 0.099 | 1.72 | 0.00 ± 0.07 | 0.957 | 2.10 |
| UV cover | 0.08 ± 0.25 | -0.305 – 0.741 | -0.250 – 0.621 | 1.13 | 0.17 ± 0.17 | -0.126 – 0.552 | -0.078 – 0.467 | 1.09 | 0.06 ± 0.07 | 0.361 | 1.19 | -0.06 ± 0.05 | 0.229 | 1.07 | 0.01 ± 0.07 | 0.891 | 1.13 | -0.05 ± 0.05 | 0.390 | 1.15 |
| Benzaldehyde | -0.34 ± 0.32 | -1.040 – 0.242 | -0.915 – 0.144 | 2.39 | 0.65 ± 0.51 | -0.273 – 1.730 | -0.110 – 1.548 | 3.15 | 0.00 ± 0.17 | 0.987 | 5.55 | -0.12 ± 0.16 | 0.483 | 6.96 | **-0.46 ± 0.12** | **<.001** | 3.48 | -0.05 ± 0.1 | 0.585 | 2.87 |
| 1-Butene-4-ITC | 0.24 ± 0.24 | -0.224 – 0.718 | -0.148 – 0.631 | 1.74 | -0.01 ± 0.24 | -0.489 – 0.445 | -0.403 – 0.366 | 1.44 | -0.09 ± 0.12 | 0.452 | 2.77 | 0.06 ± 0.09 | 0.539 | 2.27 | -0.03 ± 0.10 | 0.737 | 2.17 | *-0.16 ± 0.08* | *0.058* | 1.96 |
| Methyl benzoate | 0.01 ± 0.25 | -0.472 – 0.517 | -0.401 – 0.428 | 1.48 | 0.15 ± 0.47 | -0.697 – 1.181 | -0.561 – 0.980 | 2.08 | -0.03 ± 0.18 | 0.865 | 6.03 | 0.09 ± 0.16 | 0.594 | 6.59 | 0.17 ± 0.11 | 0.141 | 3.01 | 0.10 ± 0.10 | 0.327 | 2.97 |
| Phenylethyl alcohol | 0.76 ± 0.65 | -0.422 – 2.097 | -0.207 – 1.873 | 8.18 | 0.27 ± 0.49 | -0.760 – 1.212 | -0.568 – 1.040 | 6.52 | -0.02 ± 0.20 | 0.902 | 7.67 | -0.17 ± 0.13 | 0.197 | 4.44 | 0.04 ± 0.17 | 0.799 | 6.24 | -0.09 ± 0.11 | 0.394 | 3.43 |
| 2-Amino benzaldehyde | -0.47 ± 0.58 | -1.886 – 0.468 | -1.629 – 0.334 | 6.97 | 0.70 ± 0.61 | -0.412 – 2.036 | -0.220 – 1.751 | 5.96 | 0.09 ± 0.21 | 0.665 | 8.59 | 0.20 ± 0.14 | 0.170 | 5.25 | 0.16 ± 0.14 | 0.270 | 4.76 | -0.10 ± 0.14 | 0.485 | 6.01 |
| p-Anisaldehyde | 0.43 ± 0.36 | -0.182 – 1.267 | -0.098 – 1.116 | 3.11 | -0.12 ± 0.28 | -0.672 – 0.465 | -0.583 – 0.352 | 1.47 | 0.09 ± 0.15 | 0.515 | 4.01 | 0.06 ± 0.09 | 0.501 | 2.14 | 0.02 ± 0.13 | 0.885 | 3.56 | 0.08 ± 0.09 | 0.364 | 2.16 |
| Methyl anthranilate | 0.27 ± 0.42 | -0.656 – 1.032 | -0.487 – 0.906 | 4.62 | -0.04 ± 0.40 | -0.830 – 0.797 | -0.680 – 0.638 | 3.41 | 0.05 ± 0.18 | 0.762 | 5.94 | -0.16 ± 0.14 | 0.254 | 4.86 | -0.10 ± 0.15 | 0.506 | 5.42 | 0.07 ± 0.14 | 0.609 | 5.41 |
| 3-Hexenyl acetate | -0.01 ± 0.19 | -0.413 – 0.357 | -0.339 – 0.294 | 1.31 | -0.26 ± 0.43 | -1.139 – 0.546 | -1.000 – 0.414 | 1.93 | 0.03 ± 0.10 | 0.805 | 2.06 | -0.04 ± 0.10 | 0.695 | 2.45 | 0.15 ± 0.10 | 0.144 | 2.24 | 0.07 ± 0.08 | 0.403 | 1.75 |
| Phenylacetaldehyde | **-1.47 ± 0.75** | **-3.198 – -0.260** | -2.883 – -0.419 | 8.20 | **-1.17 ± 0.64** | **-2.692 – -0.139** | -2.435 – -0.297 | 6.51 | -0.27 ± 0.21 | 0.213 | 8.49 | 0.06 ± 0.17 | 0.728 | 7.06 | -0.17 ± 0.14 | 0.225 | 4.62 | 0.05 ± 0.09 | 0.563 | 2.37 |
| Benzyl nitrile | 0.57 ± 0.75 | -0.684 – 2.350 | -0.507 – 2.030 | 8.95 | **0.56 ± 0.33** | **0.026 – 1.355** | 0.114 – 1.195 | 3.21 | 0.04 ± 0.24 | 0.876 | 10.53 | 0.05 ± 0.14 | 0.752 | 5.39 | 0.15 ± 0.15 | 0.323 | 5.11 | 0.08 ± 0.12 | 0.508 | 4.00 |
| Methyl salicylate | 0.08 ± 0.25 | -0.451 – 0.548 | -0.357 – 0.468 | 1.83 | -0.05 ± 0.28 | -0.598 – 0.490 | -0.501 – 0.404 | 1.36 | 0.00 ± 0.18 | 0.989 | 6.43 | -0.08 ± 0.12 | 0.537 | 3.95 | -0.03 ± 0.11 | 0.778 | 2.95 | -0.10 ± 0.10 | 0.310 | 2.68 |
| Indole | -0.31 ± 0.56 | -1.456 – 0.743 | -1.251 – 0.591 | 8.81 | -0.69 ± 0.70 | -2.362 – 0.449 | -2.015 – 0.279 | 5.86 | -0.04 ± 0.27 | 0.878 | 14.22 | -0.10 ± 0.19 | 0.612 | 9.39 | -0.11 ± 0.18 | 0.532 | 7.25 | 0.01 ± 0.15 | 0.921 | 6.19 |
| *(Z.Z)*-α-Farnesene | 0.41 ± 0.69 | -0.812 – 1.867 | -0.615 – 1.609 | 10.56 | -0.60 ± 0.62 | -2.170 – 0.369 | -1.888 – 0.221 | 8.55 | -0.14 ± 0.10 | 0.181 | 2.03 | -0.11 ± 0.09 | 0.234 | 2.09 | 0.19 ± 0.17 | 0.266 | 6.75 | 0.02 ± 0.18 | 0.894 | 9.31 |
| *(E.E)*-α-Farnesene | 0.05 ± 0.66 | -1.411 – 1.160 | -1.132 – 0.978 | 9.59 | 0.58 ± 0.56 | -0.386 – 1.902 | -0.236 – 1.654 | 7.11 | **0.30 ± 0.14** | **0.040** | 3.98 | **0.23 ± 0.11** | **0.042** | 3.29 | 0.06 ± 0.16 | 0.727 | 5.79 | 0.03 ± 0.18 | 0.870 | 9.22 |

**Table S11.** The effects of temperature on the quadratic selection of plant traits under different biotic treatments. Quadratic selection gradients on plant visual traits and flower scent compounds for plants grown and tested in two temperature environments and visited by either *Bombus terrestris* bumblebees (*Bombus*), *Pieris rapae* butterflies (*Pieris*), or their combination (generalized). Visitation occurred in in an ambient (23ºC) or hot (27ºC+ weekly 30ºC day) temperature environment. where plants were previously kept for 16-18 days. Relative seed set was used as a fitness estimate in the models. Selection for morphological and scent traits were analysed separately. Because of zero-inflation in the fitness data of the *Bombus* treatment. non-parametric bootstraps were used to calculate standard errors of the gradients. and confidence intervals (C.I.) were used instead of p-values to assess significance. Bold values indicate results where *P* ≤ 0.05 or the 95% C.I. does not include zero. Italic values indicate results where *P* ≤ 0.1 or the 90% C.I. does not include zero.

| | *Bombus* | | | | | | *Pieris* | | | | Generalized | | | |
|---|---|---|---|---|---|---|---|---|---|---|---|---|---|---|
| | Ambient | | | Hot | | | Ambient | | Hot | | Ambient | | Hot | |
| | ɣ ± SE | CI (95 %) | CI (90 %) | ɣ ± SE | CI (95 %) | CI (90 %) | ɣ ± SE | P | ɣ ± SE | P | ɣ ± SE | P | ɣ ± SE | P |
| Flowering time | **-0.46 ± 0.23** | **-1.194 – -0.175** | -1.056 – -0.228 | 0.21 ± 0.29 | -0.757 – 0.416 | -0.632 – 0.302 | -0.04 ± 0.10 | 0.707 | -0.02 ± 0.04 | 0.650 | -0.09 ± 0.11 | 0.409 | 0.06 ± 0.10 | 0.520 |
| Plant height | 0.27 ± 0.46 | -0.440 – 1.468 | -0.325 – 1.241 | -0.36 ± 0.32 | -1.175 – 0.196 | -0.997 – 0.105 | 0.04 ± 0.10 | 0.724 | -0.04 ± 0.07 | 0.609 | 0.08 ± 0.11 | 0.447 | -0.10 ± 0.11 | 0.349 |
| Flower number | -0.08 ± 0.39 | -0.243 – 1.542 | -0.113 – 1.287 | 0.13 ± 0.58 | -0.611 – 1.867 | -0.485 – 1.529 | -0.03 ± 0.11 | 0.762 | -0.13 ± 0.08 | 0.122 | **-0.15 ± 0.05** | **0.003** | -0.03 ± 0.09 | 0.737 |
| Leaf number | **1.14 ± 0.43** | **0.386 – 2.215** | 0.483 – 2.018 | 0.12 ± 0.56 | -0.804 – 1.306 | -0.653 – 1.063 | **0.22 ± 0.11** | **0.046** | -0.07 ± 0.09 | 0.413 | -0.05 ± 0.09 | 0.605 | 0.05 ± 0.08 | 0.529 |
| Flower diameter | -0.02 ± 0.21 | -0.427 – 0.436 | -0.349 – 0.317 | 0.82 ± 0.62 | -0.406 – 2.131 | -0.318 – 1.798 | 0.15 ± 0.11 | 0.202 | 0.00 ± 0.10 | 0.970 | 0.08 ± 0.10 | 0.465 | 0.10 ± 0.12 | 0.377 |
| Nectar per flower | **-0.15 ± 0.16** | **-0.563 – -0.005** | -0.488 – -0.042 | **-0.69 ± 0.34** | **-1.482 – -0.085** | -1.304 – -0.180 | 0.06 ± 0.10 | 0.546 | -0.17 ± 0.09 | 0.075 | 0.14 ± 0.11 | 0.214 | -0.08 ± 0.07 | 0.260 |
| Petal area | 0.06 ± 0.28 | -0.482 – 0.745 | -0.256 – 0.616 | -0.46 ± 0.25 | -0.971 – 0.065 | -0.650 – 0.020 | -0.10 ± 0.12 | 0.442 | 0.00 ± 0.08 | 0.994 | -0.16 ± 0.12 | 0.187 | -0.05 ± 0.07 | 0.451 |
| UV cover | 0.15 ± 0.27 | -0.216 – 1.067 | -0.174 – 0.928 | *-0.38 ± 0.22* | *-0.490 – 0.007* | *-0.404 – -0.033* | -0.21 ± 0.13 | 0.100 | -0.08 ± 0.06 | 0.192 | **0.28 ± 0.08** | **0.001** | 0.02 ± 0.04 | 0.592 |
| Benzaldehyde | 0.04 ± 0.32 | -0.497 – 0.724 | -0.392 – 0.603 | -1.00 ± 0.58 | -1.621 – 0.716 | -1.414 – 0.489 | 0.06 ± 0.19 | 0.770 | 0.02 ± 0.13 | 0.856 | -0.07 ± 0.19 | 0.730 | 0.09 ± 0.11 | 0.403 |
| 1-Butene-4-isothyocyanate | -0.30 ± 0.32 | -0.961 – 0.306 | -0.830 – 0.162 | 0.01 ± 0.41 | -0.715 – 0.767 | -0.580 – 0.670 | -0.13 ± 0.16 | 0.431 | **-0.21 ± 0.11** | **0.047** | 0.09 ± 0.13 | 0.490 | -0.19 ± 0.13 | 0.157 |
| Methyl benzoate | 0.39 ± 0.34 | -0.348 – 1.125 | -0.224 – 0.921 | 0.76 ± 0.54 | -0.111 – 1.986 | -0.001 – 1.786 | 0.31 ± 0.26 | 0.238 | 0.14 ± 0.21 | 0.508 | -0.12 ± 0.14 | 0.384 | 0.07 ± 0.14 | 0.631 |
| Phenylethyl alcohol | -0.67 ± 0.40 | -0.545 – 1.086 | -0.407 – 0.900 | -0.47 ± 0.63 | -1.218 – 1.132 | -0.972 – 0.931 | 0.22 ± 0.28 | 0.439 | -0.02 ± 0.22 | 0.944 | -0.03 ± 0.16 | 0.867 | 0.01 ± 0.10 | 0.927 |
| 2-Amino benzaldehyde | -0.41 ± 0.56 | -1.876 – 0.531 | -1.513 – 0.383 | 0.40 ± 0.47 | -1.097 – 1.032 | -0.833 – 0.762 | 0.11 ± 0.21 | 0.591 | 0.20 ± 0.16 | 0.237 | -0.21 ± 0.23 | 0.357 | 0.13 ± 0.15 | 0.385 |
| p-Anisaldehyde | *-1.24 ± 0.37* | *-1.448 – 0.030* | *-1.295 – -0.096* | -0.48 ± 0.32 | -0.958 – 0.315 | -0.837 – 0.173 | -0.21 ± 0.28 | 0.457 | -0.19 ± 0.18 | 0.306 | 0.03 ± 0.18 | 0.845 | -0.17 ± 0.15 | 0.245 |
| Methyl anthranilate | -0.07 ± 0.49 | -1.443 – 0.403 | -1.264 – 0.266 | -0.39 ± 0.43 | -1.357 – 0.335 | -1.105 – 0.225 | 0.17 ± 0.25 | 0.492 | -0.07 ± 0.19 | 0.706 | 0.01 ± 0.22 | 0.966 | -0.21 ± 0.13 | 0.108 |
| *(Z)*-3-Hexen-1-ol acetate | **-0.16 ± 0.32** | **-1.268 – -0.062** | -1.099 – -0.123 | -0.08 ± 0.43 | -0.875 – 0.912 | -0.698 – 0.707 | -0.09 ± 0.15 | 0.580 | -0.14 ± 0.13 | 0.285 | -0.04 ± 0.16 | 0.784 | 0.18 ± 0.13 | 0.189 |
| Phenylacetaldehyde | 0.88 ± 0.45 | -1.199 – 0.582 | -0.992 – 0.433 | 0.85 ± 0.68 | -0.328 – 2.572 | -0.107 – 2.102 | -0.26 ± 0.31 | 0.399 | 0.28 ± 0.24 | 0.248 | -0.08 ± 0.16 | 0.633 | -0.24 ± 0.14 | 0.086 |
| Benzyl nitrile | 1.31 ± 0.88 | -0.411 – 3.226 | -0.280 – 2.610 | -0.41 ± 0.45 | -1.316 – 0.466 | -1.137 – 0.322 | 0.28 ± 0.33 | 0.397 | 0.18 ± 0.21 | 0.398 | 0.03 ± 0.20 | 0.885 | **0.31 ± 0.15** | **0.044** |
| Methyl salicylate | *-0.63 ± 0.39* | *-1.511 – 0.061* | *-1.352 – -0.049* | 0.10 ± 0.51 | -0.280 – 1.867 | -0.185 – 1.541 | -0.42 ± 0.28 | 0.141 | -0.22 ± 0.20 | 0.265 | -0.21 ± 0.17 | 0.208 | 0.21 ± 0.13 | 0.101 |
| Indole | -0.75 ± 0.45 | -1.101 – 0.718 | -0.924 – 0.516 | -0.30 ± 0.81 | -0.288 – 2.409 | -0.179 – 1.584 | -0.45 ± 0.35 | 0.199 | -0.23 ± 0.27 | 0.396 | -0.01 ± 0.19 | 0.971 | -0.10 ± 0.14 | 0.471 |
| *(Z.Z)*-α-Farnesene | **1.04 ± 0.72** | **0.101 – 3.135** | 0.258 – 2.693 | 0.57 ± 0.71 | -1.593 – 1.225 | -1.257 – 0.973 | -0.1 ± 0.18 | 0.574 | 0.07 ± 0.10 | 0.491 | 0.17 ± 0.21 | 0.429 | 0.07 ± 0.23 | 0.746 |
| *(E.E)*-α-Farnesene | -0.55 ± 0.56 | -1.904 – 0.343 | -1.639 – 0.003 | -0.67 ± 0.73 | -1.376 – 1.564 | -1.136 – 1.228 | 0.19 ± 0.20 | 0.368 | 0.14 ± 0.15 | 0.350 | -0.13 ± 0.17 | 0.427 | 0.16 ± 0.23 | 0.484 |

**Table S12.** The effects of temperature on the selection differentials of plant traits under different biotic treatments. Linear selection differentials on plant visual traits and flower scent compounds for plants grown and tested in two temperature environments and visited by either *Bombus terrestris* bumblebees (*Bombus*), *Pieris rapae* butterflies (*Pieris*), or their combination (generalized). Visitation occurred in in an ambient (23°C) or hot (27°C+ weekly 30°C day) temperature environment. where plants were previously kept for 16-18 days. Relative seed set was used as a fitness estimate in the models. Linear models regressing each trait against plant fitness were used to derive estimates. Because of zero-inflation in the fitness data of the *Bombus* treatment. non-parametric bootstraps were used to calculate standard errors of the gradients. and confidence intervals (C.I.) were used instead of p-values to assess significance. Bold values indicate results where *P* ≤ 0.05 or the 95% C.I. does not include zero. Italic values indicate results where *P* ≤ 0.1.

| | *Bombus* | | | | *Pieris* | | | | General | | | |
|---|---|---|---|---|---|---|---|---|---|---|---|---|
| | Ambient | | Hot | | Ambient | | Hot | | Ambient | | Hot | |
| | β ± SE | CI (95 %) | β ± SE | CI (95 %) | β ± SE | P | β ± SE | P | β ± SE | P | β ± SE | P |
| Flowering time | -0.01 ± 0.15 | -0.287 – 0.300 | 0.13 ± 0.22 | -0.414 – 0.500 | 0.07 ± 0.07 | 0.337 | -0.06 ± 0.06 | 0.305 | 0.08 ± 0.07 | 0.273 | -0.10 ± 0.06 | 0.098 |
| Plant height | **0.54 ± 0.24** | **0.168 – 1.148** | **0.45 ± 0.20** | **0.126 – 0.926** | **0.28 ± 0.07** | **< 0.001** | **0.23 ± 0.06** | **< 0.001** | **0.19 ± 0.07** | **0.010** | **0.17 ± 0.06** | **0.003** |
| Flower number | **0.71 ± 0.26** | **0.325 – 1.443** | 0.31 ± 0.26 | -0.076 – 1.036 | **0.32 ± 0.07** | **< 0.001** | **0.29 ± 0.05** | **< 0.001** | **0.27 ± 0.07** | **< 0.001** | **0.23 ± 0.05** | **< 0.001** |
| Leaf number | 0.46 ± 0.32 | -0.035 – 1.293 | 0.13 ± 0.24 | -0.284 – 0.646 | 0.11 ± 0.07 | 0.136 | **0.17 ± 0.06** | **0.005** | 0.03 ± 0.08 | 0.721 | **0.21 ± 0.05** | **< 0.001** |
| Flower diameter | -0.10 ± 0.21 | -0.675 – 0.228 | 0.16 ± 0.29 | -0.512 – 0.659 | 0.08 ± 0.07 | 0.249 | 0.08 ± 0.06 | 0.206 | 0.05 ± 0.07 | 0.493 | **0.12 ± 0.06** | **0.031** |
| Nectar per flower | **-0.29 ± 0.15** | **-0.662 – -0.038** | -0.05 ± 0.17 | -0.411 – 0.279 | -0.02 ± 0.07 | 0.741 | 0.05 ± 0.06 | 0.456 | -0.01 ± 0.08 | 0.888 | 0.09 ± 0.06 | 0.102 |
| Petal area | -0.24 ± 0.20 | -0.734 – 0.091 | 0.31 ± 0.19 | -0.034 – 0.708 | 0.10 ± 0.07 | 0.188 | **0.14 ± 0.06** | **0.022** | -0.01 ± 0.08 | 0.871 | 0.08 ± 0.06 | 0.151 |
| Petal length | -0.29 ± 0.26 | -0.939 – 0.127 | -0.04 ± 0.14 | -0.388 – 0.198 | 0.07 ± 0.07 | 0.356 | 0.06 ± 0.06 | 0.361 | 0.07 ± 0.07 | 0.352 | 0.05 ± 0.06 | 0.438 |
| Petal width | -0.09 ± 0.17 | -0.423 – 0.252 | 0.27 ± 0.19 | -0.074 – 0.677 | 0.12 ± 0.07 | 0.111 | **0.15 ± 0.06** | **0.011** | -0.07 ± 0.07 | 0.380 | 0.07 ± 0.06 | 0.249 |
| UV surface area | 0.19 ± 0.31 | -0.229 – 1.110 | 0.03 ± 0.15 | -0.207 – 0.362 | 0.11 ± 0.07 | 0.135 | -0.01 ± 0.06 | 0.903 | 0.02 ± 0.08 | 0.811 | -0.09 ± 0.06 | 0.115 |
| Benzaldehyde | -0.14 ± 0.20 | -0.538 – 0.245 | 0.04 ± 0.22 | -0.320 – 0.591 | 0.05 ± 0.07 | 0.474 | 0.01 ± 0.06 | 0.988 | **-0.17 ± 0.07** | **0.021** | -0.04 ± 0.06 | 0.528 |
| 1-Butene-4-ITC | 0.31 ± 0.20 | -0.008 – 0.837 | -0.03 ± 0.17 | -0.394 – 0.280 | -0.03 ± 0.07 | 0.715 | 0.03 ± 0.06 | 0.587 | -0.09 ± 0.07 | 0.257 | -0.09 ± 0.06 | 0.128 |
| Methyl benzoate | 0.02 ± 0.19 | -0.316 – 0.461 | 0.13 ± 0.31 | -0.325 – 0.975 | 0.11 ± 0.07 | 0.118 | 0.01 ± 0.06 | 0.956 | 0.01 ± 0.08 | 0.982 | 0.04 ± 0.06 | 0.538 |
| Phenylethyl alcohol | -0.13 ± 0.16 | -0.461 – 0.181 | -0.25 ± 0.17 | -0.746 – 0.001 | 0.01 ± 0.07 | 0.981 | -0.04 ± 0.06 | 0.508 | -0.05 ± 0.07 | 0.539 | -0.02 ± 0.06 | 0.793 |
| 2-Amino benzaldehyde | -0.09 ± 0.15 | -0.440 – 0.178 | 0.09 ± 0.15 | -0.214 – 0.374 | 0.08 ± 0.07 | 0.294 | 0.04 ± 0.06 | 0.540 | 0.08 ± 0.07 | 0.268 | 0.01 ± 0.06 | 0.991 |
| p-Anisaldehyde | 0.11 ± 0.17 | -0.184 – 0.486 | -0.07 ± 0.21 | -0.426 – 0.430 | 0.12 ± 0.07 | 0.112 | 0.02 ± 0.06 | 0.759 | 0.06 ± 0.07 | 0.449 | 0.06 ± 0.06 | 0.316 |
| Methyl anthranilate | 0.05 ± 0.17 | -0.273 – 0.397 | 0.10 ± 0.15 | -0.180 – 0.419 | 0.11 ± 0.07 | 0.146 | -0.02 ± 0.06 | 0.755 | 0.10 ± 0.07 | 0.162 | 0.05 ± 0.06 | 0.346 |
| *(Z)*-3-Hexen-1-ol acetate | -0.06 ± 0.13 | -0.319 – 0.180 | -0.28 ± 0.18 | -0.677 – 0.052 | 0.05 ± 0.07 | 0.456 | -0.02 ± 0.06 | 0.727 | -0.08 ± 0.07 | 0.304 | -0.01 ± 0.06 | 0.839 |
| Phenylacetaldehyde | **-0.36 ± 0.16** | **-0.666 – -0.035** | **-0.37 ± 0.20** | **-0.926 – -0.074** | -0.02 ± 0.07 | 0.832 | 0.01 ± 0.06 | 0.963 | -0.02 ± 0.08 | 0.836 | 0.02 ± 0.06 | 0.734 |
| Benzyl nitrile | 0.04 ± 0.21 | -0.296 – 0.579 | 0.05 ± 0.17 | -0.310 – 0.357 | 0.09 ± 0.07 | 0.195 | 0.01 ± 0.06 | 0.974 | 0.07 ± 0.07 | 0.357 | 0.05 ± 0.06 | 0.373 |
| Methyl salicylate | 0.10 ± 0.16 | -0.192 – 0.443 | -0.12 ± 0.23 | -0.581 – 0.317 | 0.11 ± 0.07 | 0.148 | 0.01 ± 0.06 | 0.903 | 0.01 ± 0.08 | 0.885 | 0.01 ± 0.06 | 0.983 |
| Indole | -0.01 ± 0.18 | -0.379 – 0.347 | -0.11 ± 0.19 | -0.579 – 0.187 | 0.09 ± 0.07 | 0.232 | 0.01 ± 0.06 | 0.935 | 0.03 ± 0.08 | 0.643 | 0.01 ± 0.06 | 0.844 |
| *(Z.Z)*-α-Farnesene | 0.31 ± 0.23 | -0.036 – 0.915 | -0.16 ± 0.21 | -0.639 – 0.191 | 0.02 ± 0.07 | 0.836 | -0.01 ± 0.06 | 0.927 | 0.09 ± 0.07 | 0.240 | 0.03 ± 0.06 | 0.590 |
| *(E.E)*-α-Farnesene | 0.29 ± 0.21 | -0.021 – 0.882 | -0.04 ± 0.21 | -0.550 – 0.314 | **0.15 ± 0.07** | **0.035** | 0.06 ± 0.06 | 0.353 | 0.09 ± 0.07 | 0.208 | 0.02 ± 0.06 | 0.796 |

**Table S13.** The directional selection of temperature-mediated plasticity under different biotic treatments. Directional selection gradients on the temperature reaction norms of plant visual traits and flower scent compounds for sibling plants visited by either *Bombus terrestris* bumblebees (*Bombus*), *Pieris rapae* butterflies (*Pieris*), or their combination (generalized). Reaction norms were calculated comparing trait values of full-sibling plants grown in an ambient (23°C) and hot (27°C+ weekly 30°C day) temperature environment. Total family relative seed set was used as a fitness estimate in the models. Selection for morphological and scent traits were analysed separately. Because of zero-inflation in the fitness data of the *Bombus* treatment. non-parametric bootstraps were used to calculate standard errors of the gradients. and confidence intervals (C.I.) were used to assess significance. Bold values indicate results where $P \leq 0.05$ or the 95% C.I. does not include zero. Italic values indicate results where $P \leq 0.1$ or the 90% C.I. does not include zero. Scent emission is expressed as pg / flower / l / h.

| | *Bombus* | | | | *Pieris* | | | Generalized | | |
|---|---|---|---|---|---|---|---|---|---|---|
| | β ± SE | CI (95 %) | CI (90 %) | VIF | β ± SE | P | VIF | β ± SE | P | VIF |
| Flowering time | 0.22 ± 0.16 | -0.071 – 0.569 | -0.025 – 0.502 | 1.07 | 0.05 ± 0.05 | 0.286 | 1.05 | -0.03 ± 0.06 | 0.638 | 1.12 |
| Plant height | *0.33 ± 0.23* | *-0.045 – 0.877* | *0.011 – 0.770* | 1.59 | 0.03 ± 0.06 | 0.566 | 1.54 | 0.01 ± 0.06 | 0.809 | 1.25 |
| Flower number | -0.32 ± 0.23 | -0.814 – 0.108 | -0.732 – 0.044 | 1.29 | -0.01 ± 0.05 | 0.871 | 1.30 | -0.04 ± 0.06 | 0.548 | 1.17 |
| Leaf number | -0.18 ± 0.21 | -0.659 – 0.195 | -0.577 – 0.135 | 1.45 | 0.01 ± 0.05 | 0.779 | 1.15 | **0.13 ± 0.06** | **0.025** | 1.14 |
| Flower diameter | -0.04 ± 0.21 | -0.423 – 0.413 | -0.352 – 0.338 | 1.63 | -0.09 ± 0.06 | 0.151 | 1.70 | 0.03 ± 0.07 | 0.649 | 1.92 |
| Nectar per flower | -0.16 ± 0.16 | -0.481 – 0.154 | -0.424 – 0.100 | 1.18 | **0.14 ± 0.05** | **0.009** | 1.26 | 0.05 ± 0.06 | 0.364 | 1.07 |
| Petal area | 0.22 ± 0.23 | -0.159 – 0.748 | -0.096 – 0.649 | 1.78 | 0.02 ± 0.06 | 0.753 | 1.66 | 0.01 ± 0.07 | 0.948 | 1.77 |
| UV cover | -0.16 ± 0.18 | -0.579 – 0.148 | -0.505 – 0.100 | 1.16 | -0.08 ± 0.05 | 0.122 | 1.10 | -0.08 ± 0.05 | 0.145 | 1.14 |
| Benzaldehyde | **0.66 ± 0.28** | **0.141 – 1.256** | 0.220 – 1.154 | 2.47 | -0.05 ± 0.10 | 0.615 | 4.69 | 0.05 ± 0.08 | 0.519 | 2.58 |
| 1-Butene-4-isothyocyanate | 0.02 ± 0.18 | -0.341 – 0.375 | -0.274 – 0.306 | 1.80 | 0.12 ± 0.07 | 0.126 | 2.52 | 0.02 ± 0.07 | 0.828 | 2.01 |
| Methyl benzoate | -0.03 ± 0.20 | -0.419 – 0.376 | -0.345 – 0.299 | 1.67 | -0.05 ± 0.12 | 0.684 | 6.82 | -0.06 ± 0.08 | 0.451 | 2.85 |
| Phenylethyl alcohol | *-0.72 ± 0.44* | *-1.653 – 0.062* | *-1.496 – -0.055* | 5.65 | -0.01 ± 0.11 | 0.907 | 5.14 | 0.01 ± 0.09 | 0.966 | 3.49 |
| 2-Amino benzaldehyde | 0.14 ± 0.43 | -0.703 – 0.980 | -0.568 – 0.819 | 8.63 | 0.12 ± 0.12 | 0.340 | 6.80 | -0.11 ± 0.13 | 0.410 | 7.03 |
| p-Anisaldehyde | -0.43 ± 0.32 | -1.003 – 0.261 | -0.917 – 0.131 | 2.48 | *-0.14 ± 0.08* | *0.086* | 3.11 | -0.02 ± 0.07 | 0.801 | 2.15 |
| Methyl anthranilate | 0.03 ± 0.29 | -0.546 – 0.607 | -0.433 – 0.512 | 3.86 | -0.16 ± 0.11 | 0.175 | 5.81 | 0.05 ± 0.11 | 0.666 | 5.13 |
| *(Z)*-3-Hexen-1-ol acetate | -0.02 ± 0.19 | -0.421 – 0.336 | -0.348 – 0.274 | 1.47 | -0.11 ± 0.07 | 0.113 | 2.03 | 0.03 ± 0.07 | 0.673 | 2.15 |
| Phenylacetaldehyde | **0.89 ± 0.48** | **0.065 – 1.918** | 0.162 – 1.724 | 5.61 | 0.11 ± 0.11 | 0.316 | 5.62 | 0.02 ± 0.09 | 0.792 | 3.00 |
| Benzyl nitrile | -0.30 ± 0.42 | -1.185 – 0.484 | -1.030 – 0.348 | 4.81 | -0.10 ± 0.14 | 0.498 | 9.09 | 0.10 ± 0.08 | 0.213 | 2.80 |
| Methyl salicylate | -0.24 ± 0.22 | -0.795 – 0.112 | -0.690 – 0.062 | 1.51 | 0.01 ± 0.12 | 0.993 | 5.99 | 0.09 ± 0.08 | 0.251 | 2.41 |
| Indole | 0.56 ± 0.50 | -0.407 – 1.576 | -0.245 – 1.373 | 10.13 | 0.12 ± 0.14 | 0.385 | 8.86 | 0.01 ± 0.13 | 0.970 | 6.62 |
| *(Z.Z)*-α-Farnesene | -0.71 ± 0.52 | -1.759 – 0.300 | -1.584 – 0.138 | 7.74 | -0.02 ± 0.06 | 0.797 | 1.63 | 0.08 ± 0.1 | 0.447 | 4.28 |
| *(E.E)*-α-Farnesene | 0.32 ± 0.46 | -0.570 – 1.264 | -0.414 – 1.097 | 6.75 | 0.12 ± 0.08 | 0.170 | 3.15 | -0.17 ± 0.11 | 0.125 | 4.75 |

**Table S14.** The non-linear selection of temperature-mediated plasticity under different biotic treatments. Quadratic selection gradients on the temperature reaction norms of plant visual traits and flower scent compounds for sibling plants visited by either *Bombus terrestris* bumblebees (*Bombus*), *Pieris rapae* butterflies (*Pieris*), or their combination (generalized). Reaction norms were calculated comparing trait values of full-sibling plants grown in an ambient (23°C) and hot (27°C+ weekly 30°C day) temperature environment. Total family relative seed set was used as a fitness estimate in the models. Selection for morphological and scent traits was analysed separately. Because of zero-inflation in the fitness data of the *Bombus* treatment. non-parametric bootstraps were used to calculate standard errors of the gradients. and confidence intervals (C.I.) were used to assess significance. Bold values indicate results where $P \leq 0.05$ or the 95% C.I. does not include zero. Italic values indicate results where $P \leq 0.1$ or the 90% C.I. does not include zero.

| | *Bombus* | | | *Pieris* | | Generalized | |
|---|---|---|---|---|---|---|---|
| | β ± SE | CI (95 %) | CI (90 %) | β ± SE | P | β ± SE | P |
| Flowering time | -0.14 ± 0.17 | -0.522 – 0.170 | -0.450 – 0.088 | 0.09 ± 0.07 | 0.188 | -0.15 ± 0.12 | 0.217 |
| Plant height | -0.33 ± 0.22 | -0.632 – 0.264 | -0.549 – 0.156 | -0.02 ± 0.07 | 0.723 | 0.01 ± 0.07 | 0.987 |
| Flower number | 0.56 ± 0.33 | -0.147 – 1.166 | -0.019 – 1.069 | -0.02 ± 0.07 | 0.735 | -0.02 ± 0.06 | 0.785 |
| Leaf number | 0.20 ± 0.39 | -0.285 – 1.313 | -0.182 – 1.141 | 0.06 ± 0.07 | 0.357 | **0.16 ± 0.07** | **0.036** |
| Flower diameter | 0.18 ± 0.29 | -0.388 – 0.751 | -0.298 – 0.604 | 0.01 ± 0.08 | 0.852 | *0.20 ± 0.11* | *0.065* |
| Nectar per flower | 0.11 ± 0.26 | -0.505 – 0.458 | -0.444 – 0.390 | **0.16 ± 0.07** | **0.030** | -0.07 ± 0.09 | 0.429 |
| Petal area | -0.10 ± 0.25 | -0.417 – 0.551 | -0.338 – 0.226 | -0.01 ± 0.08 | 0.899 | -0.15 ± 0.09 | 0.091 |
| Petal length | 0.03 ± 0.33 | -0.433 – 0.623 | -0.347 – 0.540 | *-0.13 ± 0.07* | *0.057* | 0.03 ± 0.05 | 0.490 |
| UV cover | -0.22 ± 0.40 | -0.942 – 0.640 | -0.846 – 0.412 | 0.13 ± 0.09 | 0.187 | -0.03 ± 0.09 | 0.715 |
| Benzaldehyde | -0.01 ± 0.32 | -0.317 – 1.023 | -0.235 – 0.832 | **-0.25 ± 0.10** | **0.018** | *0.18 ± 0.09* | *0.057* |
| 1-Butene-4-isothyocyanate | -0.12 ± 0.29 | -0.978 – 0.109 | -0.828 – 0.023 | 0.07 ± 0.15 | 0.613 | -0.03 ± 0.07 | 0.688 |
| Methyl benzoate | 0.08 ± 0.49 | -0.597 – 1.268 | -0.485 – 1.081 | -0.22 ± 0.13 | 0.092 | -0.11 ± 0.10 | 0.241 |
| Phenylethyl alcohol | 0.26 ± 0.49 | -1.098 – 0.818 | -0.909 – 0.653 | -0.06 ± 0.14 | 0.693 | 0.06 ± 0.15 | 0.661 |
| 2-Amino benzaldehyde | -0.57 ± 0.43 | -1.085 – 0.659 | -0.962 – 0.449 | -0.23 ± 0.14 | 0.110 | -0.07 ± 0.07 | 0.302 |
| p-Anisaldehyde | 0.01 ± 0.39 | -0.663 – 0.895 | -0.532 – 0.743 | 0.16 ± 0.17 | 0.362 | **-0.25 ± 0.10** | **0.022** |
| Methyl anthranilate | -0.39 ± 0.36 | -1.178 – 0.236 | -1.001 – 0.146 | -0.01 ± 0.10 | 0.939 | -0.02 ± 0.10 | 0.848 |
| *(Z)*-3-Hexen-1-ol acetate | -0.03 ± 0.32 | -0.833 – 0.384 | -0.688 – 0.283 | 0.05 ± 0.16 | 0.754 | 0.02 ± 0.11 | 0.885 |
| Phenylacetaldehyde | -0.21 ± 0.51 | -1.154 – 0.877 | -1.011 – 0.662 | 0.06 ± 0.17 | 0.734 | -0.12 ± 0.11 | 0.302 |
| Benzyl nitrile | -0.08 ± 0.29 | -0.659 – 0.482 | -0.571 – 0.352 | 0.07 ± 0.16 | 0.677 | -0.04 ± 0.12 | 0.738 |
| Methyl salicylate | 0.19 ± 0.44 | -0.579 – 1.183 | -0.401 – 0.990 | -0.16 ± 0.19 | 0.393 | 0.12 ± 0.16 | 0.451 |
| Indole | 0.62 ± 0.60 | -0.188 – 2.173 | -0.021 – 1.971 | 0.16 ± 0.10 | 0.130 | -0.20 ± 0.13 | 0.120 |
| *(Z.Z)*-α-Farnesene | -0.31 ± 0.49 | -1.544 – 0.409 | -1.335 – 0.204 | -0.06 ± 0.09 | 0.551 | -0.09 ± 0.09 | 0.344 |
| *(E.E)*-α-Farnesene | -0.14 ± 0.17 | -0.522 – 0.170 | -0.450 – 0.088 | 0.09 ± 0.07 | 0.188 | -0.15 ± 0.12 | 0.217 |

**Table S15.** Effects of temperature on patterns of additive and nonadditive linear selection. Estimated additive and observed linear selection gradients for plants visited by bumblebees and butterflies in an ambient (23°C) and hot (27°C+ weekly 30°C day) temperature environment. Additive selection gradients were calculated by summing the selection gradients of plants pollinated by either bumblebees or butterflies alone. Nonadditive?: A significant interaction ($P \leq 0.05$) between *data* and *flower trait* for the combined predicted and observed dataset indicates nonadditive selection. Separate models were run for visual and scent traits. Bold values indicate results where $P \leq 0.05$. Italic values indicate results where $P \leq 0.1$.

| | Ambient | | | | Hot | | | |
|---|---|---|---|---|---|---|---|---|
| | Additive | Observed | | Nonadditive? | Additive | Observed | | Nonadditive? |
| | $\beta_{ADD}$ ± SE | $\beta_{O}$ ± SE | P | P | $\beta_{ADD}$ ± SE | $\beta_{O}$ ± SE | P | P |
| Flowering time | 0.11 ± 0.25 | 0.10 ± 0.08 | 0.254 | 0.866 | -0.03 ± 0.27 | -0.07 ± 0.05 | 0.178 | **<0.001** |
| Plant height | 0.61 ± 0.33 | *0.15 ± 0.08* | *0.052* | **<0.001** | 0.48 ± 0.26 | 0.08 ± 0.06 | 0.187 | **<0.001** |
| Flower number | 0.94 ± 0.33 | **0.25 ± 0.07** | **0.001** | **<0.001** | 0.66 ± 0.32 | *0.10 ± 0.06* | *0.083* | **<0.001** |
| Leaf number | -0.20 ± 0.36 | -0.03 ± 0.07 | 0.691 | **0.017** | 0.33 ± 0.35 | **0.19 ± 0.06** | **0.001** | **<0.001** |
| Flower diameter | -0.15 ± 0.37 | 0.12 ± 0.09 | 0.176 | **0.002** | -0.15 ± 0.46 | 0.09 ± 0.07 | 0.233 | **0.001** |
| Nectar per flower | -0.41 ± 0.23 | -0.07 ± 0.07 | 0.321 | **<0.001** | -0.13 ± 0.31 | 0.04 ± 0.05 | 0.450 | **<0.001** |
| Petal area | 0.07 ± 0.33 | -0.16 ± 0.09 | 0.099 | **0.017** | 0.51 ± 0.33 | 0.00 ± 0.07 | 0.957 | 0.359 |
| UV surface area | 0.14 ± 0.32 | 0.01 ± 0.07 | 0.891 | *0.073* | 0.11 ± 0.22 | -0.05 ± 0.05 | 0.390 | **<0.001** |
| Benzaldehyde | -0.34 ± 0.49 | **-0.46 ± 0.12** | **<0.001** | **<0.001** | 0.53 ± 0.67 | -0.05 ± 0.1 | 0.585 | 0.584 |
| 1-Butene-4-ITC | 0.15 ± 0.36 | -0.03 ± 0.10 | 0.737 | *0.063* | 0.05 ± 0.33 | -0.16 ± 0.08 | 0.058 | **<0.001** |
| Methyl benzoate | -0.02 ± 0.43 | 0.17 ± 0.11 | 0.141 | *0.099* | 0.24 ± 0.63 | 0.10 ± 0.10 | 0.327 | 0.239 |
| Phenylethyl alcohol | 0.74 ± 0.85 | 0.04 ± 0.17 | 0.799 | **<0.001** | 0.10 ± 0.62 | -0.09 ± 0.11 | 0.394 | **<0.001** |
| 2-Amino benzaldehyde | -0.38 ± 0.79 | 0.16 ± 0.14 | 0.270 | **<0.001** | 0.90 ± 0.75 | -0.10 ± 0.14 | 0.485 | *0.056* |
| p-Anisaldehyde | 0.52 ± 0.51 | 0.02 ± 0.13 | 0.885 | **<0.001** | -0.06 ± 0.37 | 0.08 ± 0.09 | 0.364 | **<0.001** |
| Methyl anthranilate | 0.32 ± 0.6 | -0.10 ± 0.15 | 0.506 | **0.007** | -0.20 ± 0.54 | 0.07 ± 0.14 | 0.609 | *0.071* |
| *(Z)*-3-Hexen-1-ol acetate | 0.02 ± 0.29 | 0.15 ± 0.10 | 0.144 | 0.205 | -0.30 ± 0.53 | 0.07 ± 0.08 | 0.403 | 0.559 |
| Phenylacetaldehyde | -1.74 ± 0.96 | -0.17 ± 0.14 | 0.225 | **<0.001** | -1.11 ± 0.81 | 0.05 ± 0.09 | 0.563 | **<0.001** |
| Benzyl nitrile | 0.61 ± 0.99 | 0.15 ± 0.15 | 0.323 | **0.002** | 0.61 ± 0.47 | 0.08 ± 0.12 | 0.508 | **<0.001** |
| Methyl salicylate | 0.08 ± 0.43 | -0.03 ± 0.11 | 0.778 | 0.326 | -0.13 ± 0.4 | -0.10 ± 0.10 | 0.310 | *0.066* |
| Indole | -0.35 ± 0.83 | -0.11 ± 0.18 | 0.532 | 0.185 | -0.79 ± 0.89 | 0.01 ± 0.15 | 0.921 | **0.014** |
| *(Z.Z)*-α-Farnesene | 0.27 ± 0.79 | 0.19 ± 0.17 | 0.266 | 0.655 | -0.71 ± 0.71 | 0.02 ± 0.18 | 0.894 | 0.174 |
| *(E.E)*-α-Farnesene | 0.35 ± 0.8 | 0.06 ± 0.16 | 0.727 | *0.067* | 0.81 ± 0.67 | 0.03 ± 0.18 | 0.870 | *0.076* |

**Table S16.** Effects of temperature on patterns of additive and nonadditive selection on temperature-mediated plasticity. Estimated additive and observed selection gradients on the temperature reaction norms of plants visited by bumblebees and butterflies in an ambient (23°C) and hot (27°C+ weekly 30°C day) temperature environment. Additive selection gradients were calculated by summing the selection gradients of plants pollinated by either bumblebees or butterflies alone. Nonadditive?: A significant interaction ($P \leq 0.05$) between *data* and *flower trait* for the combined predicted and observed dataset indicates nonadditive selection. Separate models were run for visual and scent traits. Bold values indicate results where $P \leq 0.05$. Italic values indicate results where $P \leq 0.1$. Traits with missing values were excluded from multiple regression selection analyses due to collinearity.

| | Linear | | | |
|---|---|---|---|---|
| | Predicted | Observed | | Nonadditive? |
| | $\beta_{ADD}$ ± SE | $\beta_O$ ± SE | P | P |
| Flowering time | 0.27 ± 0.21 | -0.03 ± 0.06 | 0.638 | **<0.001** |
| Plant height | 0.36 ± 0.29 | 0.01 ± 0.06 | 0.809 | **<0.001** |
| Flower number | -0.33 ± 0.28 | -0.04 ± 0.06 | 0.548 | **<0.001** |
| Leaf number | -0.17 ± 0.26 | **0.13 ± 0.06** | **0.025** | **<0.001** |
| Flower diameter | -0.13 ± 0.27 | 0.03 ± 0.07 | 0.649 | **0.023** |
| Nectar per flower | -0.02 ± 0.21 | 0.05 ± 0.06 | 0.364 | 0.205 |
| Petal area | 0.24 ± 0.29 | 0.01 ± 0.07 | 0.948 | **<0.001** |
| UV surface area | -0.24 ± 0.23 | -0.08 ± 0.05 | 0.145 | **0.004** |
| Benzaldehyde | 0.61 ± 0.38 | 0.05 ± 0.08 | 0.519 | **<0.001** |
| 1-Butene-4-ITC | 0.14 ± 0.25 | 0.02 ± 0.07 | 0.828 | *0.079* |
| Methyl benzoate | -0.08 ± 0.32 | -0.06 ± 0.08 | 0.451 | 0.844 |
| Phenylethyl alcohol | -0.73 ± 0.55 | 0.01 ± 0.09 | 0.966 | **<0.001** |
| 2-Amino benzaldehyde | 0.26 ± 0.55 | -0.11 ± 0.13 | 0.410 | **0.006** |
| p-Anisaldehyde | -0.57 ± 0.4 | -0.02 ± 0.07 | 0.801 | **<0.001** |
| Methyl anthranilate | -0.13 ± 0.4 | 0.05 ± 0.11 | 0.666 | 0.114 |
| *(Z)*-3-Hexen-1-ol acetate | -0.13 ± 0.26 | 0.03 ± 0.07 | 0.673 | **0.029** |
| Phenylacetaldehyde | 1.00 ± 0.59 | 0.02 ± 0.09 | 0.792 | **<0.001** |
| Benzyl nitrile | -0.40 ± 0.56 | 0.10 ± 0.08 | 0.213 | **<0.001** |
| Methyl salicylate | -0.23 ± 0.34 | 0.09 ± 0.08 | 0.251 | **<0.001** |
| Indole | 0.68 ± 0.64 | 0.01 ± 0.13 | 0.970 | **<0.001** |
| *(Z.Z)*-α-Farnesene | -0.73 ± 0.58 | 0.08 ± 0.10 | 0.447 | **<0.001** |
| *(E.E)*-α-Farnesene | 0.44 ± 0.54 | -0.17 ± 0.11 | 0.125 | **<0.001** |
